# Hybrid spatial organization and magnitude-independent neural coding of linguistic information during sentence production

**DOI:** 10.1101/2024.06.20.599931

**Authors:** Adam M. Morgan, Orrin Devinsky, Werner K. Doyle, Patricia Dugan, Daniel Friedman, Adeen Flinker

## Abstract

Humans are the only species with the ability to systematically combine words to convey an unbounded number of complex meanings. This process is guided by combinatorial processes whose underlying neural mechanisms remain obscured by inherent limitations of non-invasive brain measures and a near total focus on comprehension paradigms. Here, we address these limitations with high-resolution neurosurgical recordings (electrocorticography) and a controlled sentence production experiment. We uncover distinct cortical networks encoding word-level and higher-order information. These networks exhibited a hybrid spatial organization: broadly distributed across traditional language areas, but with focal concentrations of sensitivity to semantic and structural contrasts in canonical language regions. In contrast to previous comprehension-based findings, we find that these networks are largely non-overlapping. Most strikingly, our data reveal an unexpected property of higher-order linguistic information: it is encoded independent of neural activity levels. These results show that activity magnitude and information content are dissociable, with important implications for studying the neurobiology of language.

**Teaser:** “Brain recordings during speech reveal complex linguistic information throughout cortex, independent of neural activity levels.”

---

Sentence production is the process by which the brain translates from thought to language. Despite its centrality in human cognition, daily life, and in a host of debilitating neurological disorders, it remains poorly characterized in the brain due to a number of critical gaps in the literature. For one, previous research has relied primarily on non-invasive neuroimaging and electrophysiology (e.g., fMRI, EEG, MEG), methods which make overt production very difficult to study given their susceptibility to motion and/or motor artifacts. Furthermore, these methods impose a trade-off between spatial and temporal resolution (*1, 2*), both of which are needed to study a dynamic, anatomically distributed system like language. Another (perhaps related) gap is that the neuroscience of language has nearly exclusively focused on either sentence comprehension or single-word production (*3, 4*), with only a handful of studies to date investigating the production of multi-word utterances (*5–18*).

One of the complexities of studying sentences is that, relative to single words, they involve higher-order constructs like combinatorial semantics and syntax. While these systems are rarely investigated with production, they have been extensively studied in the comprehension literature. However, far from elucidating, this body of work contains a number of discrepant findings, and significant disagreements persist regarding the basic facts. One such area of disagreement is *localization*: how these systems are spatially encoded. For semantic composition – the process of combining the meanings like “red” and “boat” into a single representation of a “red boat” – some research points toward a hub in left anterior temporal lobe (*19–23*), while other research points toward a broader network spanning left lateral cortex (*7, 24, 25*). For syntax, there is similarly little clarity. Some research has traced syntax back to inferior frontal gyrus (IFG) (*6, 26–36*) with other work identifying posterior temporal areas (*18, 26–33, 37–39*). A recent influential neural model of syntax in the brain (*33*) assigns functions to the various regions associated with syntax. This model holds that posterior temporal areas are responsible for forming hierarchically structured representations of a sentence, IFG (particularly pars triangularis) is responsible for transforming these into linear strings of words (or, more specifically, morphemes), and that this is then translated into a sequence of phonemes for articulation by posterior IFG (pars opercularis), dorsal pre-central gyrus, and the temporo-parietal junction (*33*). However, ROI-based models like this face a challenge in accounting for a growing body of findings indicating that, rather than being localized to particular regions, syntax is distributed across broad swaths of cortex (*34, 39–42*).

Relatedly, there is significant disagreement regarding *selectivity*: the degree to which these systems overlap spatially, which may have important theoretical implications (see (*42, 43*) for discussion). While findings of syntax-selective regions are abundant in the literature (*29–31, 39, 44–51*), recent work using more targeted analytical techniques has reported extensive spatial overlap in these systems (*39–42, 52–54*), which has been argued to be evidence that there may in fact be no areas of the brain that process syntax to the exclusion of words and/or meaning (*42*). However, other work involving highly sensitive neural measures indicates that spatial overlap and functional selectivity may be able to co-exist, enabled by distinct microcircuits in the same region, particularly IFG (*36*).

At least three factors have contributed to these discrepancies and disagreements. First, different studies define syntax differently, and may in fact be studying different phenomena. Despite often being treated as a monolith, syntax is in fact a complex system involving multiple types of representations (e.g., function words, hierarchical “tree” structures, word categories) and processes (e.g., agreement, sequencing, and binding) (*30, 33, 55–57*). Some experimental designs target the depth of syntactic processing by manipulating the complexity of hierarchical structures (*27, 34, 40*), while others have manipulated the presence or absence of syntactic structure by, for instance, comparing sentence stimuli to unstructured word lists (*27,52,54,58–64*), aiming to target the system as a whole. Indeed, one emerging pattern is that grosser comparisons like this tend to identify larger networks, potentially highlighting their utility as a localizer for higher-order linguistic processing, but also highlighting the need for finer-grained experimental comparisons to delineate finer-grained anatomical distinctions. For example, findings of broad spatial distributions are common for sentence-list comparisons [e.g., (*52*)], but finer-grained dissociations of hierarchical vs. linear properties (*37*) have resulted in the kind of regional-functional specificity outlined above [e.g., (*33*)].

The second factor relates the difficulty of isolating a system of abstract representations. To experimentally activate abstract syntactic representations – e.g., a “rule” like *Sentence = Subject + Verb + Object* – one must use words, which often introduces confounds with properties like phonology and meaning, complicating interpretation (*3, 55*). For instance, in what has become one of the most common paradigms in this area, sentences are compared to unstructured lists of words (*27, 52, 54, 58–64*). The underlying assumption – a pervasive assumption that we will return to throughout this manuscript – is that elevated neural activity indicates information processing. For convenience, we will refer to this as the *Elevation-Processing Equivalence* assumption (or **EPE**). The EPE has a long tradition dating back to early sensory neuroscience, where classical studies demonstrated a direct correspondence between the intensity of sensory stimuli and the rate of firing of sensory neurons [e.g., (*65, 66*)]. Formal information–theoretic treatments similarly treat firing rate as a carrier of information (*67, 68*). Within such a framework, it is natural to assume that the EPE holds for all types of information processing – from low-level sensory to higher-order cognitive. More recently, functional imaging and electrophysiology approaches have reinforced this view: correlates of neuronal firing (e.g., alpha/beta suppression in M/EEG, BOLD signal in fMRI, high gamma power in intracranial EEG) are standardly interpreted as transparent indices of information processing [e.g., (*69–71*)]. On this assumption, a brain region that shows higher activity for sentences than lists would be taken as processing information that is present in sentences but not lists. However, sentences and lists differ not only the presence of syntax and combinatorial semantics, but a number of other factors including depth of processing, ecological validity, prosody, and so on, making it difficult to uniquely attribute differences to any one particular construct. Another common approach is to vary particular syntactic structures while holding words (*19, 38*) and/or meaning (*12, 72, 73*) constant. For instance, active and passive sentences can be used to express the same meaning by arranging the same content words (e.g., Dracula, spray, chicken) in reverse orders (active: “Dracula sprayed the chicken”; passive: “The chicken was sprayed by Dracula”). This varies structure while holding constant meaning, lexical content, and sentence processing. However, comparing actives and passives comes with its own set of confounds: active and passive sentences differ in processing difficulty, and prosodic features, structural frequency, and the number and type of function words they involve (e.g., the passive *be* and *by*).

The final factor that contributes to discrepancies in the literature is the historical reliance on comprehension paradigms. Comprehension studies offer some clear benefits: they provide a high degree of experimental control, allowing the researcher to present a precise stimulus at a precise time. And to be sure, comprehension is a useful tool for studying language in general, as higher-order linguistic representations are shared across the two modalities (*12, 74, 75*). However, relative to production, comprehension introduces a number of modality-specific complications. For one, it engages the language network to a lesser degree than production (*13*), rendering signal detection more difficult. Another (perhaps related) complication is that syntax is often ignored during comprehension in favor of less effortful strategies (e.g., context) (*76–79*). In production, the act of successfully producing a grammatical sentence entails syntactic processing, mitigating such concerns. Finally, comprehension is an ongoing process of building and assessing candidate structures and meanings in response to a continually changing stream of input (*29, 57, 80, 81*) – a highly dynamic process that renders reasonable temporal granularity difficult to achieve, particularly when attempting to examine specific representations (*12, 14, 38, 82*). In contrast, in production critical higher-order representations like the global structure and meaning of a sentence are selected by the time production begins and remain invariant over time, providing a short, clearly defined time window – the *planning period* – during which to study these representations (*7, 8, 13, 29, 81, 83, 84*).

Here, we address these gaps with a controlled sentence production experiment and by sampling neural data directly from cortex in ten neurosurgical patients. Electrocorticography (ECoG), which is virtually immune to motor artifacts, provides simultaneously high spatial and temporal resolution, allowing us to precisely characterize neural activity across left lateral cortex during sentence production on a millisecond scale. Beyond just mapping activity, we aimed to characterize the spatiotemporal dynamics of higher-order linguistic processing. To avoid the types of confounds discussed above (and mitigate the risk of false positives associated with encoding models (*85, 86*)), we combined two complementary contrasts: a sentence>list test and an active≠passive test. Consistent with the EPE assumption, the sentence>list contrast was intended to capture higher-order processes engaged during sentence production, such as combinatorial semantics and syntactic processing, while controlling for articulation, multi-word production, and word-level processes, which are all shared by both sentences and lists. Importantly, sentences and lists may also differ in domain-general processes such as working memory and attention, although it is not clear a priori in which direction such differences should go, if at all. The active≠passive contrast also targets syntactic structure, along with a *different* set of nuisance properties, including the presence of particular words (e.g., the passive ‘by’), prosody, and the engagement of domain-general control processes like attention and working memory. Rather than assuming that either contrast isolates “syntax” in a pure sense, we focused on their intersection, reasoning that because the two contrasts both isolate syntax in common but little else, regions sensitive to both contrasts would be more likely to reflect syntax – and more specifically, representations of particular syntactic structures (or derivational processes, depending on the particular syntactic theory [see (*87*) for discussion]; we remain agnostic here) – while acknowledging that contributions from domain-general functions may not be fully eliminated by either comparison alone. Critically, this intersection was intended as an initial operationalization of syntactic processing under the EPE. However, as we will show below, the results of this analysis motivate a re-evaluation of the basic assumptions of the EPE and a more fine-grained analysis of the information encoded at individual electrodes. Together, the combination of ECoG, a controlled production paradigm, and two tests for higher-order linguistic processing afford unparalleled resolution.

## 1 Results

Ten neurosurgical patients with electrode coverage of left peri-Sylvian cortex (Fig. 1A) performed a sentence production task and two control tasks: list production and picture naming (Fig. 1B). During sentence production, patients overtly described cartoon vignettes depicting transitive actions (e.g., poke, scare, etc.) in response to a preceding question. Questions were manipulated to use either active (“Who poked whom?”) or passive syntax (“Who was poked by whom?”), implicitly priming patients to respond with the same structure (“The chicken poked Dracula” or “Dracula was poked by the chicken”) (*88*). This manipulation held semantic and lexical content largely constant while varying syntactic structure and associated constructs like the number of function words, prosodic properties, and cognitive effort. During the list production control task, participants saw the same vignettes as in the sentence production trials, but preceded by an arrow indicating the direction in which participants should list the characters: left-to-right (e.g., “chicken Dracula”) or right-to-left (“Dracula chicken”), mimicking the reverse order of nouns in active and passive sentences. We quantified neural activity as high gamma broadband activity (70 − 150 Hz), normalized (*z*-scored) to each trial’s 200 ms pre-stimulus baseline, as this correlates with underlying spiking and BOLD signal (*89, 90*) and has been widely employed in the ECoG literature. Parallel analyses on beta-band activity, also associated with cognition (*91, 92*), appear in the supplemental information (see Supplemental Information 3.10).

**Figure 1:**
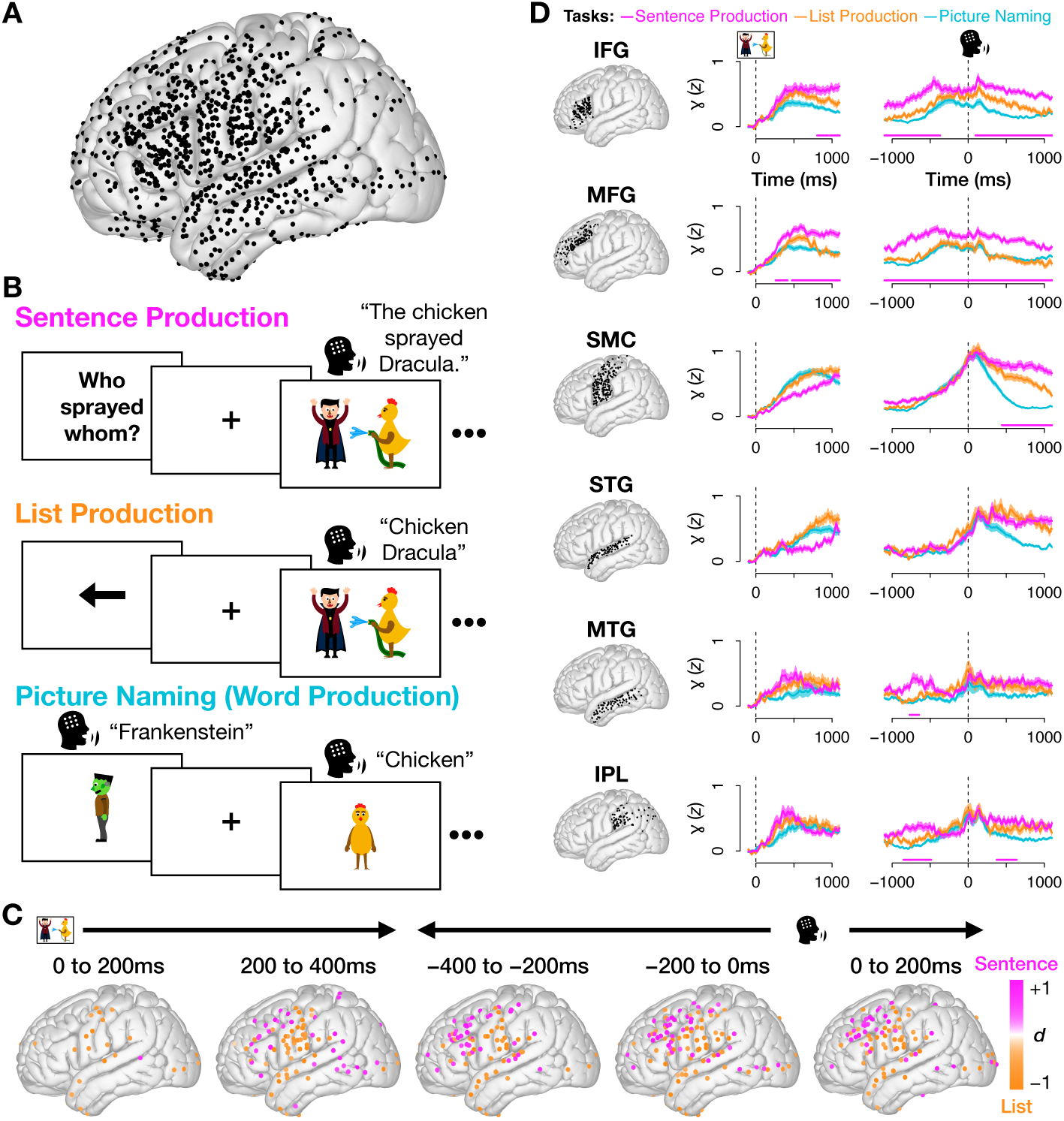
Methods and neural activity during overt language production. (A) Coverage across all 10 patients (1256 electrodes after exclusions). (B) Experimental design: Participants completed three overt production blocks. In Sentence Production, they produced sentences to respond to preceding questions. Questions appeared with either active (“Who sprayed whom?”) or passive syntax (“Who was sprayed by whom?”), implicitly priming corresponding responses (active: “The chicken sprayed Dracula”; passive: “Dracula was sprayed by the chicken”). In List Production, a control task, participants saw the Sentence Production stimuli again, but preceded by an arrow indicating the order to name two characters in. In Picture Naming, they produced single words. (N.B.: The cartoon characters shown here, created by the authors, are not those used in the experiment.) (C) Magnitude of difference in neural activity (Cohen’s *d*) for all electrodes with a significant difference between sentences and lists, in 200 ms bins. (D) Mean and standard error of neural activity for each task, by region, locked to picture onset (left) and speech onset (right). Pink bar (bottom) indicates times when sentences had significantly greater activity than lists. **Region abbreviations:** sensorimotor cortex (SMC); inferior parietal lobe (IPL); gyri: inferior frontal (IFG); middle frontal (MFG); superior temporal (STG); middle temporal (MTG).

We began by comparing neural activity between sentence production and the list production control task, a contrast commonly used to isolate higher-order linguistic processing. We looked for differences specifically during the planning period – the time between the onset of the cartoon vignette and speech onset, when higher-order features of the sentence like syntax, event semantics, and prosody are planned (*7, 8, 29, 81, 84, 93*). (In our stimuli, this is logically ensured by the fact that the first word of the sentence – the subject – depends on the choice of syntactic structure, making it impossible to begin speaking without a priori structure assignment.) This test identified 60 broadly distributed electrodes with significantly higher activity for sentences (*p* < 0.05 for 100 ms, Wilcoxon rank sum test). Consistent with previous findings, sentences recruited higher activity than lists in the 200 to 400 ms post-stimulus window in posterior temporal areas and the inferior parietal lobule (IPL), and increasing in IFG and middle frontal gyrus (MFG) leading up to production (Fig. 1C shows the distribution of all electrodes with significant differences between sentences and lists over time; *p* < 0.05, FDR-corrected across electrodes, Wilcoxon rank sum test). A region-of-interest (ROI) analysis (Fig. 1D) found four regions with significantly higher activity for sentences during the planning period: IFG, MFG, IPL, and middle temporal gyrus (MTG) (*p* < 0.05 for at least 100 ms; one-tailed Wilcoxon rank sum test; see Methods Sec. 3.7) – all regions commonly associated with higher-order language processing (*3,33,57*). These results are nicely complemented by the inverse effects in the beta band analyses (Fig. S4), where suppression is generally interpreted as indexing cognitive processing (*94, 95*). This was particularly true in sensorimotor cortex, which showed higher activity for sentences than lists in the beta band (Fig. S4C), the mirror of what was observed for high gamma. Interestingly, this difference manifested less clearly in associative cortex. This likely reflects known spatial heterogeneity in the relationship between beta and local activation: in sensorimotor (and other lower-level) cortices, beta decreases are often tightly coupled to increases in high-frequency activity and are therefore commonly interpreted as a local index of processing, whereas in associative/cognitive-control networks beta can reflect more distributed, network-level dynamics (e.g., top–down control or state maintenance) whose relationship to local high gamma is less consistent (*94–97*).

While sentence>list tests are common, these behaviors differ in multiple processes (*54, 58, 59, 61–63*). We therefore aimed to better isolate higher-order linguistic information by intersecting these results with those of another comparison: active vs. passive sentences, which differ in syntactic structure, number of words (the passive *be* and *by*), and the difficulty of grammatical encoding. Passive sentences had longer initiation latencies than actives (see Sec. 3.8), likely reflecting this difficulty. To remove these differences, we followed (*98–100*) and temporally warped the data, setting each trial’s response time to the median for that task (see Supplementary Fig. S1). Such temporal warping has previously been shown to increase signal-to-noise ratio, a finding which we successfully replicated in the current dataset (Supplementary Fig. S3). Supplementary Fig. S2 shows the results of applying the same analyses from Fig. 1 to the warped data; the pattern of significant results remains unchanged. Figure 2A shows the sentence>list and active≠passive tests for three sample electrodes. Electrodes E1 and E2 showed patterns we expected to find based on previous work: E1 had significantly higher activity for sentences than lists (top plot; *p* < 0.05 for 100 ms, Wilcoxon rank sum test) and corresponding differences between active and passive syntax in the same time windows (bottom; *p* < 0.05 for 100 ms, Wilcoxon rank sum test). Electrode E2 also had significantly higher sentence activity (*p* < 0.05 for 100 ms, Wilcoxon rank sum test), but showed no sensitivity to structure, suggesting involvement in some other process that differs between sentences and lists such as combinatorial semantics.

**Figure 2:**
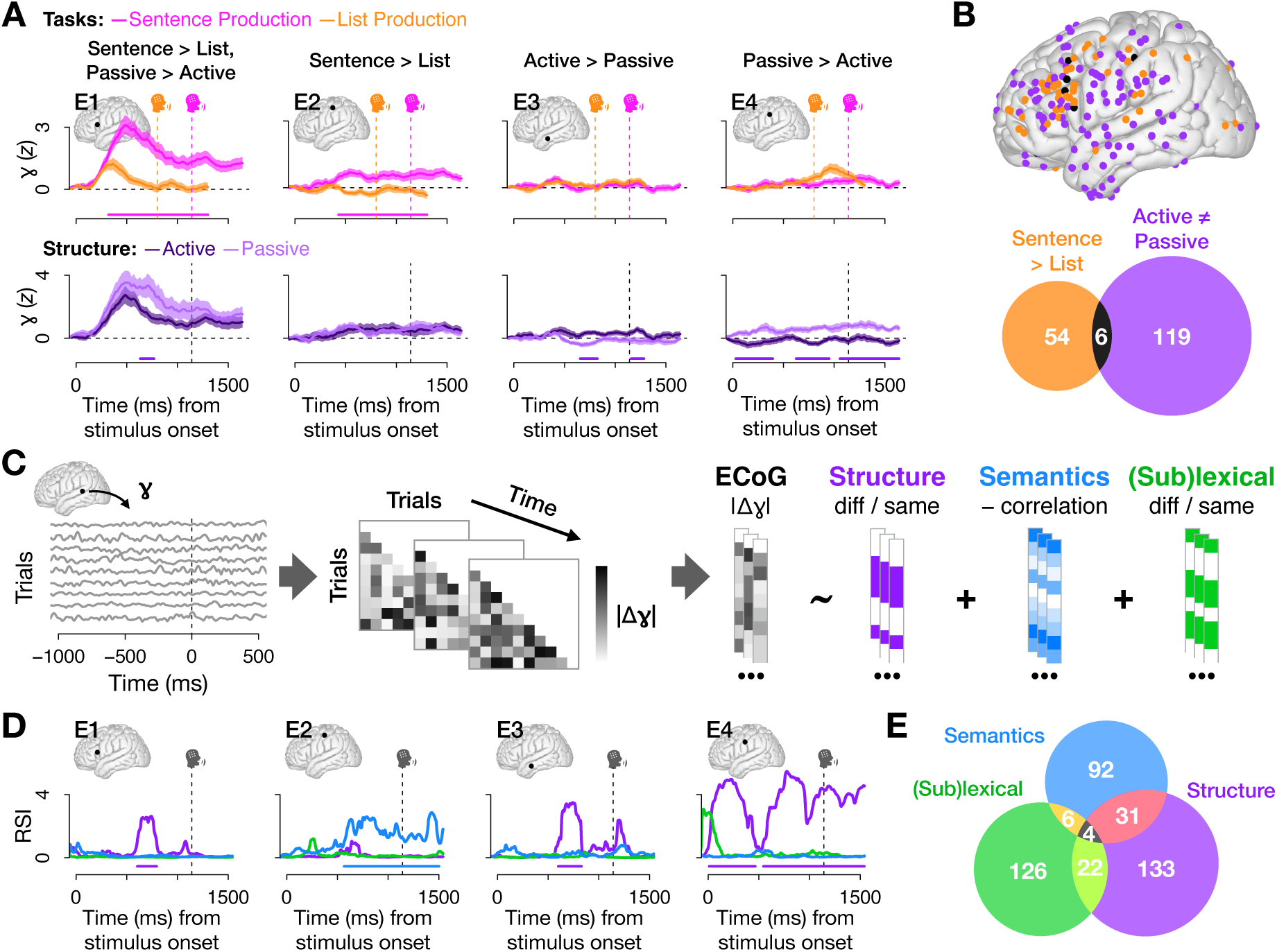
Comparing two approaches to identifying higher-order linguistic information. (A) Four sample electrodes. Top: mean and standard error of high gamma activity by task. Pink bars denote where sentences were significantly higher than lists (*p* << 0. 0.05 for 100 ms, Wilcoxon rank sum test). Bottom: mean and standard error of sentence trials split by sentence structure; bars denote significant differences between active vs. passive trials (< 0.*p* < 0.05 for 100 ms, Wilcoxon rank sum test). (B) 60 electrodes had significantly higher activity for sentences than list< 0.05 for 100 ms, Wilcoxon rank sum test) and 125 electrodes had significant differences between active and passive trials (*p* < 0.05 for 100 ms, Wilcoxon rank sum test). Only 6 electrodes (< 5) had both. (C) RSA analysis pipelines (*p* < 0.05 for 100 ms, Wilcoxon rank sum test) and 125 electrodes had significant differences between active and passive trials (*p* < 0.05 for 100 ms, Wilcoxon rank sum test). Only 6 electrodes (< 5) had both. (C) RSA analysis pipeline: For each electrode at each time sample, we modeled the magnitude of differences in high gamma activity for each pair of sentence trials as a function of differences in structure (active/passive), event semantics, and (sub)lexi< 0.05 for 100 ms; one-tailed permutation tests). (E) Significant electrodes (*p* < 0.05 for 100 ms, permutation test) by RSA term. Most significant electrodes were selective for just one model term>cal content. (D) *Representational Similarity Indices* (RSIs; derived from model coefficients) for the same four sample electrodes show sensitivity to the active/passive contrast in E1, E3, and E4 and event semantics in E2 (all *p* < 0.05 for 100 ms; one-tailed permutation tests). (E) Significant electrodes (*p* < 0.05 for 100 ms, permutation test) by RSA term. Most significant electrodes were selective for just one model term.

However, the most prevalent pattern we observed among significant electrodes, exemplified by Electrodes E3 and E4, was unexpected. Despite no significant increase in sentence activity relative to lists, these electrodes showed significant differences between active and passive trials. These differences could in principle be due to any feature that differs between actives and passives, including syntactic structure, difficulty, utterance length, or prosody. However, all of these features are involved to a higher degree in sentences than lists, and we would therefore expect that they should all be processed in regions with higher activity during sentence than list production. The combination of sensitivity to syntactic frame without elevated sentence activity therefore seems to violate a common assumption in the field: that information processing corresponds to increased neural activity. Indeed, while 125 electrodes were identified by the active vs. passive, only 6 of these (fewer than 5%) were identified by comparing sentences to lists (Fig. 2B). Thus, whatever properties are isolated in the active/passive comparison are not well characterized by “increased neural activity.” This lack of overlap calls into question the EPE assumption, providing evidence that information processing and elevated activity are dissociable: the active/passive contrast reveals robust sentence-level information in many cortical sites that do not exhibit increased activity relative to list production. Notably, this small set of overlapping electrodes appears to have some spatial organization. Of the six electrodes detected by both contrasts, five were located in prefrontal cortex. This spatial concentration suggests that the limited overlap may reflect domain-general executive processes such as working memory, response selection, or motor planning, which one can reasonably assume are differentially engaged by both contrasts and are known to be localized to prefrontal cortex. With broader prefrontal coverage or a larger sample, it is possible that this spatial organization would emerge more clearly; however, the fact that it remains rare even with substantial prefrontal sampling suggests that such processes account for only a small fraction of sentence-level information encoding. Independent of localization, the striking lack of overlap further indicates that the initial intersection-based approach, while conceptually well-motivated, is insufficient to capture the full range of sentence-level information encoded in cortex. We therefore next turn to an analysis that directly quantifies the information present at individual electrodes, independent of overall activity magnitude.

To quantify and isolate sources of variance, we leveraged an analytical technique that combines Representational Similarity Analysis (RSA) and multiple regression (Fig. 2C; Methods Sec. 3.9) (*101–104*). This approach allows us to more cleanly identify what kinds of information particular electrodes encode without relying on overall activity levels. We modeled neural dissimilarity as a linear combination of event-semantic (GPT-2 sentence embeddings (*105*)), structural (active vs. passive), (sub)lexical (same vs. different sentence-initial word), and response time dissimilarity. The (sub)lexical term accounts for variance associated with surface differences between active and passive sentences such as the presence of the word “by” – although these differences did not appear until the third or fourth word of the sentences (relatively late compared to the period prior to sentence onset, where we concentrated our analyses). Similarly, the structure term encompasses not only the syntactic structure of the utterance, but also the prosody, number of function words, and encoding difficulty; we use the word “structure” to refer to this term (rather than, e.g., “syntax”) to remain agnostic as to which (combination) of these properties drive effects related to this distinction. We also included response time as a covariate in the model to further account for potential differences in the frequency and/or difficulty of actives and passives. We derived *Representational Similarity Indices* (RSIs) from the model coefficients to approximate the amount of evidence for structure-related, event semantic, and (sub)lexical information in each electrode (shown for the four sample electrodes in Fig. 2D). The structure RSI accurately captured the presence of differences related to the structure of the sentence in E1, E3, and E4, and verified that E2’s higher activity for sentences than lists corresponded to the presence of event semantics (*p* < 0.05 for 100 ms, permutation test, for all three tests). Across electrodes with any significant RSIs, the majority of electrodes processed only one of the three types of information (Fig. 2E), contradicting claims of a fully interwoven system (*41, 52, 53*) but consistent with an overlapping microcircuitry framework (*36*) (see the anatomical distribution of these areas of overlap in Fig. S7).

The puzzling lack of correlation between degree of neural activity and sensitivity to the active/passive comparison observed in Electrodes E3 and E4 motivated us to further investigate this relationship. We performed a series of analyses comparing each RSI to high gamma. Note that while RSIs index differences in high gamma activity in an electrode, such differences do not require that the electrode exhibit a high overall degree of activity. This, however, is the general assumption, and predicts that RSIs should be positively correlated with the magnitude of high gamma activity. In two analyses we found no evidence that either event semantics or the various differences targeted in the active/passive comparison were related to neural activity in this way: not across electrodes at the time when each electrode’s RSI reaches its maximum (Fig. 3A), nor across electrodes at each time sample (Fig. 3B). In contrast, (sub)lexical processing showed the expected result in both analyses: a positive relationship with degree of high gamma activity at the times when the (sub)lexical RSI peaked (*p* < 0.001, linear regression), as well as at each time sample (*p* < 0.05 for 100 ms, Spearman correlation). Thus, intriguingly, the independence from the magnitude of neural activity appears to correspond specifically to the two higher-order linguistic terms.

**Figure 3:**
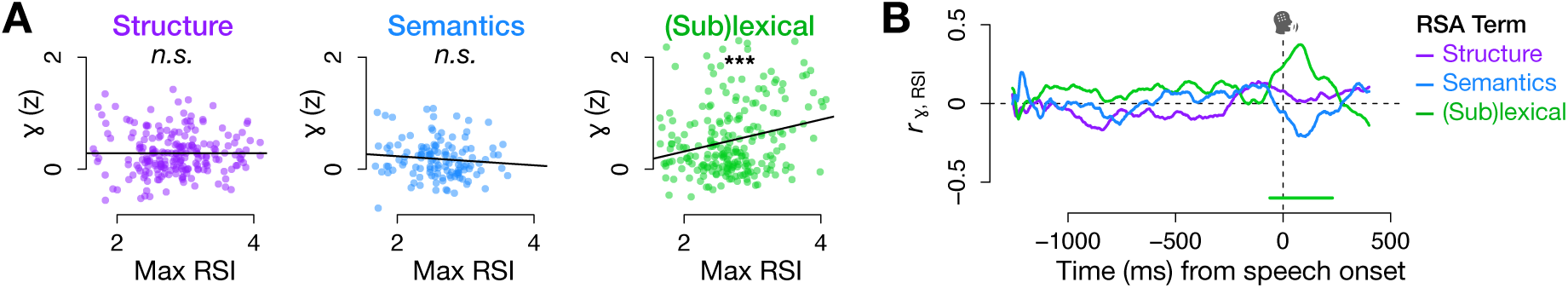
Representational Similarity Indices reveal higher-order linguistic coding decoupled from high gamma magnitude. This figure depicts two approaches to assessing the relationship between high gamma activity and RSIs for significant electrodes. In both, (sub)lexical processing was significantly correlated with the degree of neural activity, while no such evidence was found for either of the two higher-order terms. (A) Regressing activity over RSIs at the electrode- and RSI-specific time sample when RSIs peaked (structure: *p* = 0.987, semantics: *p* = 0.248, (sub)lexical: *p* < 0.001, linear regression). (B) Correlating across electrodes at each time sample (significant (sub)lexical RSI: *p* < 0.05 for 100 ms, Spearman correlation).

A traditional region-of-interest approach (Fig. 4A) showed above-chance concentrations of sensitivity to the active/passive contrast in IFG and MFG, event-semantics in MTG and IPL, and (sub)lexical information in SMC and STG (*p* < 0.05 for 100 ms, permutation test). However, none of these three types of information were limited to particular regions. Within-electrode testing (Fig. 4B) revealed that these terms were widely distributed across cortex. (See also Supplementary Fig. S6 for the same tests on beta-band activity, which similarly showed a broad distribution across cortex. At the level of ROIs, we found significant structure sensitivity in IFG and SMC but not MFG (Fig. S6A), despite robust structure effects in MFG in high gamma, consistent with our earlier observation that beta–high-gamma coupling is less consistent in associative/domain general cognitive control areas (*95–97*).) Given that our findings were not confined to particular cortical hubs, we aimed to characterize these networks without imposing classical anatomical assumptions. We took a data-driven approach and used an unsupervised machine learning technique, Non-negative Matrix Factorization (NMF; see Methods 3.10), to cluster electrodes according to prototypical patterns in the combined high gamma and RSI dataset (Fig. 4C). This identified five major networks of electrodes (Fig. 4D).

**Figure 4:**
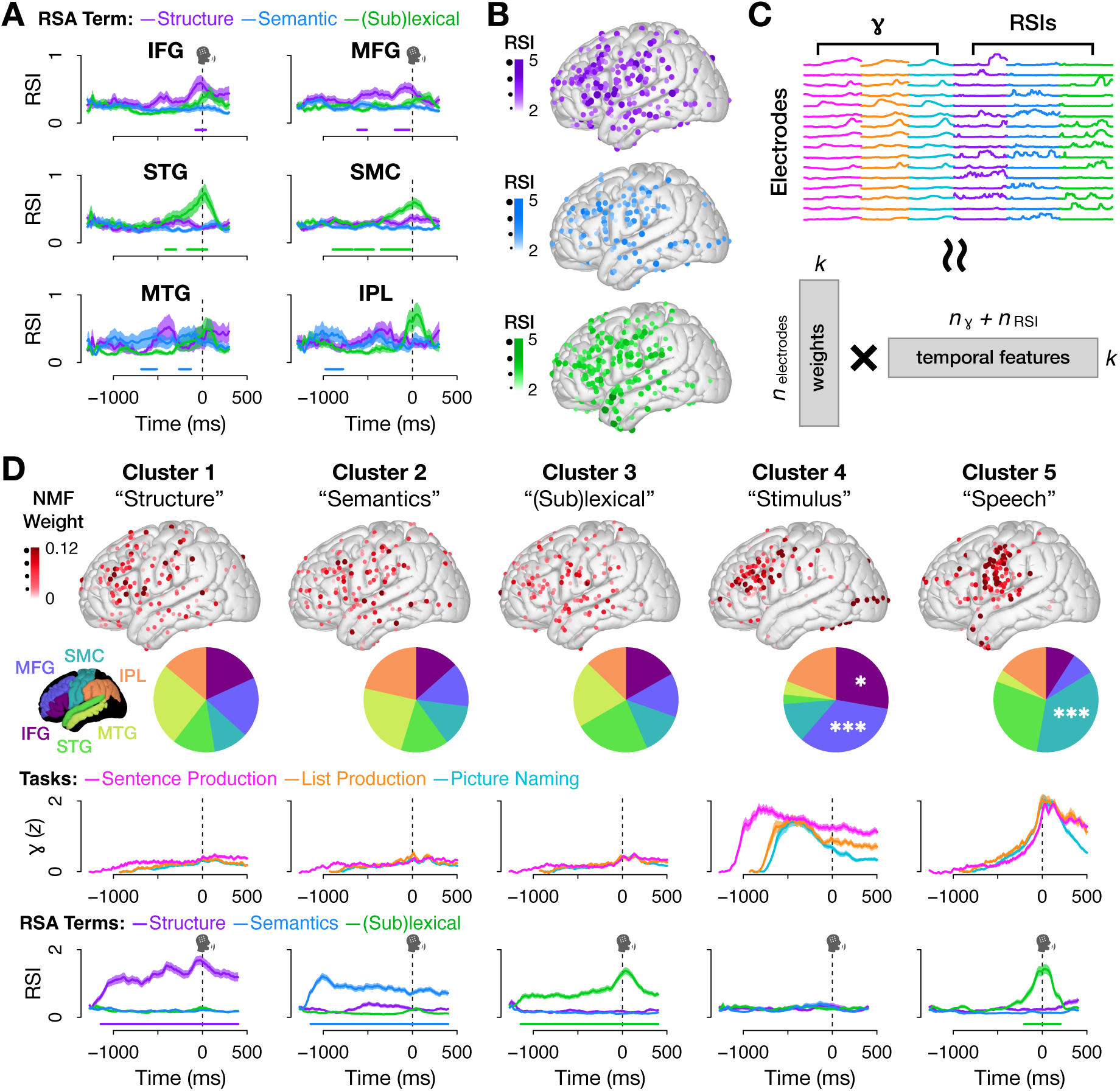
Low-activity linguistic codes and high-activity stimulus/response networks emerge from unsupervised clustering. (A) Mean and standard error of electrodes’ RSIs by ROI. (B) Significant electrodes (*p* < 0.05 for 100 ms; one-tailed permutation test) for all three RSIs; color and size correspond to peak RSI value. (C) Non-negative Matrix Factorization (NMF), a matrix decomposition technique well-suited for identifying prototypical patterns, was used to cluster electrodes based on their concatenated high gamma activity (trial means for the three tasks) and RSIs. (D) Clusters. Top: electrode localizations; shade and size correspond to NMF weight. Pie charts show ROI distributions weighted by overall electrode coverage (ROI color key appears to the left). Asterisks denote ROIs significantly over-represented in a cluster relative to chance. Middle: weighted mean and standard error of high gamma for all three experimental tasks. Bottom: weighted mean and standard error of RSIs. Bars denote significance (*p* < 0.001 for 100 ms, permutation test).

The first three clusters were characterized by high information content (RSI) accompanied by very low high gamma activity (Clusters 1-3 in Fig. 4D), replicating the dissociation between structure-dependent information and neural activity that we observed at the electrode-level (Fig. 2B). Each of these three clusters was spatially distributed, showing no above-chance electrode counts in any of our six ROIs (pie charts), and contained above-chance levels of information for just one model term – event semantics, structure, or (sub)lexical information (*p* < 0.05 for 100 ms, permutation test). These clusters demonstrate that high information content does not require elevated activity magnitude at the level of distributed cortical networks.

In contrast, the last two clusters were characterized by robust high gamma activity and high regional specificity, despite the fact that our unsupervised approach did not have access to electrode locations. Cluster 4 was focused in areas associated with executive functioning like attention and working memory (IFG, *p* = 0.039; MFG, *p* < 0.001; Fisher’s exact test, FDR-corrected across ROIs), and additionally had a number of electrodes in areas associated with visual processing (occipital lobe, although this was not included among our ROIs as it is not a traditional language region). Cluster 4’s high gamma activity peaked just after stimulus onset, and it did not contain any above-chance linguistic information (*p* > .05, permutation test across all time points after correction for multiple comparisons). Cluster 5 was concentrated in regions associated with articulation and sensory feedback (i.e., SMC, *p* < 0.001; Fisher’s exact test, FDR-corrected across ROIs), with activity peaking just after speech onset. It contained above-chance information only for the (sub)lexical RSI (*p* < 0.05 for 100 ms, permutation test) at the time when activity peaked, consistent with the correlation between (sub)lexical processing and neural activity (Fig. 3). The presence of this latter cluster serves as an internal validation of the NMF approach: despite jointly clustering high gamma activity and RSIs in a single, undifferentiated feature space, the algorithm groups these measures together when joint patterns exist (as for sensorimotor and (sub)lexical processing), but separates them for higher-order linguistic information.

## 2 Discussion

Here, we leveraged the high spatial and temporal resolution of ECoG to investigate the dynamics of sentence production. Adopting the Elevation-Processing Equivalence – the assumption that elevated activity indexes information processing – we performed a sentence>list test, a standard approach to separating (sub)lexical information from higher-order constructs like syntax and combinatorial semantics. This comparison identified a number of broadly distributed electrodes, but with focal concentrations in commonly identified regions: IFG, MFG, MTG, and IPL (Fig. 1C,D). An active≠passive test similarly identified a broadly distributed network, with focal concentrations in IFG and MFG. Given that both of these analyses were meant to identify higher-order linguistic information, we expected that identifying the electrodes significant for both should highlight higher-order language with particularly high precision, removing false positives and electrodes engaged in processes not targeted by both contrasts. Instead, the near absence of overlap between these contrasts highlights a dissociation between neural processing and elevated activity: sentence-level information can be encoded, even in the absence of activity increases. Traditional region-of-interest (ROI) analyses appeared to validate this logic, identifying regions frequently associated with higher-order language like IFG and MFG. However, ECoG provides the spatial resolution to look at smaller patches of cortex – those covered by single electrodes. Among the electrodes that showed significant sensitivity to the active/passive contrast, fewer than 5% were also significant for the sentence>list test (Fig. 2B).

An important consideration in interpreting both the sentence>list and active≠passive contrasts is the potential contribution of domain-general processes such as working memory, attention, and cognitive control. These processes are likely more engaged during passives than actives, and possibly differentially engaged for sentences and lists (although in which direction is not a priori clear). Our design mitigates this concern to some extent, as producing lists, active sentences, and passive sentences all require multi-word production, sustained attention, and maintenance of upcoming lexical items during a planning period. Moreover, domain-general demands alone would be expected to modulate overall activity magnitude, whereas we observe that higher-order information related to sentence structure and event semantics is frequently present without corresponding increases in high-gamma activity. Nonetheless, we cannot fully rule out contributions from domain-general functions, and we therefore interpret both contrasts as reflecting mixtures of linguistic and non-linguistic processes rather than pure indices of, e.g., syntax or combinatorial semantics. Critically, however, because of the near total lack of overlap between the sentence>list and active≠passive contrasts, we can conclude that magnitude independence is a property of any processes subsumed under the active≠passive contrast.

On the hypothesis that this discrepancy might reflect the coarseness of these comparisons, both of which target a wide array of sentence-level language processes, we used an RSA-based approach to separate sources of variance in individual electrodes (Fig 2C–E). From the model results, we derived “Representational Similarity Indices” (RSIs), which characterize the degree of sensitivity to event-semantic information, (sub)lexical information, and remaining information sensitive to the active/passive distinction. In a traditional ROI-based approach, we found some regional specificity for each of the three RSIs, with MTG and IPL showing above-chance sensitivity to event semantics, IFG and MFG to the active/passive contrast, and SMC and STG to (sub)lexical information (Fig. 4B). However, as before, this only partially captured the distribution: there were significant electrodes for each of the three RSIs in every region we looked at.

This suggests a hybrid organization – networks that are broadly distributed, but with focal concentrations. To characterize these networks without imposing classical anatomical assumptions, we performed an unsupervised clustering analysis (NMF) on the joint pattern of neural activity (high gamma) and information processing (RSIs) (Fig. 4C), separating electrodes into five distinct networks (Fig. 4D). Two of these were characterized by peaks in neural activity and, despite the clustering algorithm not having access to electrodes’ localizations, high spatial concentrations reflecting well-established regional-functional links: between visual processing and occipital lobe, attention and prefrontal cortex, articulation and motor cortex, and auditory feedback and STG. The remaining three clusters were defined functionally rather than by activity patterns: each uniquely encoded information related to one of the three RSIs. Intriguingly, these functional clusters recapitulated the finding of magnitude independence – originally shown at the electrode level (Fig. 3) – at the network level, as, despite high information content, they were all characterized by overall low activity.

These results contribute to an emerging picture of language in the brain as more nuanced than initially thought. Previous research has resulted in a number of discrepancies, particularly regarding localization and selectivity. Our results are consistent with previous findings of a broadly distributed code (*7, 24, 25, 34, 39–42*) and also with focal hubs (*6, 18–23, 26–38*), indicating that these two possibilities are not mutually exclusive, as they are often treated. A hybrid spatial code would not only reconcile localization discrepancies, but may also explain disagreements regarding selectivity. For instance, previous ECoG work on sentence comprehension identified spatially overlapping microcircuits in IFG that were differentially sensitive to syntactic and semantic violations (*36*). Our finding of selectivity for different types of higher-order linguistic information in the same regions is consistent with this claim, and extends it beyond just IFG to left peri-Sylvian cortex more broadly.

### 2.1 Magnitude-Independent Coding and the EPE

A widespread assumption that dates back to at early sensory neuroscience (*65, 66*) holds that information processing corresponds to areas with elevated neural activity – what we have referred to as the EPE. However, we found that higher-order linguistic information was uncorrelated with the degree of neural activity measured. That is, while many electrodes showed differences in activity corresponding to higher-order linguistic distinctions, those differences did not systematically correspond to times when the overall degree of activity in the electrode was high. This “magnitude-independent coding” scheme may help account for the lack of overlap between electrodes identified by the sentence>list and active≠passive tests, as, of the two, only the sentence>list test relies on the EPE being true. If it is not, there would need to be considerable adjustments made to tests the field uses to target higher-order language.

The striking dissociation between the presence of higher-order linguistic information and degree of neural activity may not be so surprising in light of other findings. For instance, the well-established phenomenon of repetition suppression (*106–109*), whereby the amount of neural activity involved in processing a particular representation decreases with repetition, necessitates a certain degree of independence between information processing and the overall degree of neural activity. Furthermore, multivariate analytical approaches (e.g., multivariate pattern analysis, or MVPA) are commonly employed in cognitive neuroscience precisely because of the increased sensitivity they offer for detecting relatively small differences in activity, which indicates that, even if tacitly, researchers have collectively begun to recognize that the EPE may not always be reliable. Indeed, even for some lower-level systems like olfaction and vision, neural codes outside a simple rate/population code have been observed that would not predict a strict coupling between information processing and elevated activity (*110, 111*).

Several neuroscientific studies of language comprehension studies have similarly employed RSA representational geometry to study language organization rather than exclusively focusing on overall response magnitude. This work has relied on comprehension data from non-invasive measures. Results have shown, for instance, that IFG resolves ambiguities in lexico-syntactic information (*112*), lexico-semantic encoding in posterior middle temporal lobe (*112, 113*), and modality-independent semantic sensitivity in the left intraparietal sulcus (*113*). More recent work has similarly leveraged fine-grained differences in neural activity patterns corresponding to particular sentences to reveal similar representational architectures in artificial neural network language models (*114*). (Intriguingly, this latter study used an activity magnitude-based localizer to restrict fMRI voxels within which to perform RSA, which our findings indicate may have inadvertently excluded important regions for encoding fine-grained linguistic detail.) Together, these comprehension studies demonstrate the utility of fine-grained analytical approaches like RSA for looking at information content. Our study extend this approach to sentence production in intracranial recordings. Furthermore, by directly comparing an RSA-style analysis with activity magnitude contrasts, the present work reveals an important dissociation between the two.

Regarding this claim of magnitude independence, a couple of conceptual and methodological points bear emphasis. Conceptually, it is vital to distinguish between the degree of activity (“elevation”) on one hand, and small differences in activity, which can (and do, as we have shown) occur in areas regardless of their overall activity magnitude. We argue that what is relevant is not the small differences, but the fact that the cortical sites where differences occurred did not systematically exhibit globally elevated activity. Methodologically, it is important to note that even though our claim of magnitude-independent coding is based on comparing the magnitude of high-gamma activity with representational similarity indices (RSIs) that are themselves derived from the same high-gamma activity, there is no statistical circularity here. Crucially, these two measures capture different aspects of the signal. Mean activity is a summary of the analytic amplitude, which captures the magnitude of the signal. The RSIs on the other hand are based on trial-wise differences activity, which effectively subtract out magnitude. In practice, for each electrode and time sample we computed a model fit on the trial-wise representational dissimilarity matrix and then converted this statistic into an RSI by z-scoring it relative to a null distribution obtained by shuffling trial labels. This non-parametric procedure avoids distributional assumptions about analytic amplitude. It also down-weights potential outliers (a risk due to residual ictal activity), which are equally expressed in the shuffled null. As a result, a site can exhibit a strong RSI (i.e., carry information about structure or event semantics) regardless of its mean high-gamma activity levels, and our magnitude-independence claim rests on a comparison between two conceptually distinct summaries of the same signal, rather than on re-using the same statistic in a circular way.

If higher-order linguistic information does involve a magnitude-independent code, it would require reassessing certain common assumptions about the relationship between information processing and degree of neural activity. It would also have important methodological implications: revealing a risk in relying on elevated activity as a test for the presence of information; cautioning against the common practice of a priori excluding electrodes/voxels/regions with low-activity responses [e.g., (*85, 85, 105, 114–119*)]; and underscoring the importance of fine-grained analytical tools (e.g., RSA and MVPA). In terms of experimental design, an important consequence would be that sentence-list comparisons should be interpreted with caution, perhaps particularly in production. That is because the sentence>list test relies on the assumption that higher-order linguistic processing elevates neural activity. However, in our data this identified only a small fraction of electrodes sensitive to a structural manipulation, which should have homed in on many of the same cognitive constructs (Fig. 2B). The current experiment does not allow us to definitively determine which of the many differences between actives and passives (e.g., word count, difficulty, syntactic structure) drove significant effects in this comparison. However, this may not be highly consequential: the near total absence among the active≠passive electrodes of sentence>list electrodes indicates that in general, structure-dependent production processes like prosodic planning and grammatical encoding do not elevate activity more than list production.

As for why this might be the case, there are a number of possibilities. It may be related to the nature of production, in which case our findings would not necessarily contradict those of previous comprehension studies. One reasonable hypothesis is that list processing is fundamentally different between comprehension and production. Comprehension being more passive, it may be possible for participants to “turn off” costly higher-order processes such as attempting to identify a licit syntactic parse or extracting complex meaning from the input (*52, 76, 77*). In production, on the other hand, lists may *elevate* activity with respect to sentences, as list production is more contrived and less practiced, potentially making it more difficult. If so, then it is to be expected that sentence>list comparisons would provide different results in comprehension and production. Another possibility is simply that, despite the prevalence of the sentence>list test in the literature, list comprehension may not control for syntax as intended. Previous research shows that single words engage syntactic processing (*120–122*) and that comprehending apparently ungrammatical sequences of words triggers syntactic analysis (*123–125*). Both of these findings suggest that list comprehension might also engage syntax. Moreover, the subjective ability to recognize a list is not a sentence seems to logically entail syntactic processing. Given this, it may even be reasonable to hypothesize that lists engage syntax *more* than sentences, as the comprehension system struggles to identify a syntactic structure that fits the input. Taken together, these points suggest that more work is needed to fully understand how lists recruit language networks in comprehension and production, and indicate a need for finer-grained manipulations (see (*56*) for discussion).

### 2.2 Localization: A hybrid spatial organization

Across analyses we observed a hybrid spatial organization: effects were broadly distributed across traditional language regions (Figs. 1C & 4B), but also showed focal concentrations – in IFG and MFG for the structure RSI; STG and SMC for the (sub)lexical RSI; and MTG and IPL for the event semantic RSI (Fig. 4A). (Note that regional concentrations were assessed using permutation tests that preserve the original electrode distribution across regions, ensuring that focal effects reflect above-chance clustering rather than heterogeneous sampling density.) This hybrid pattern could help reconcile ongoing debates, particularly in the syntax literature where there remains significant disagreement regarding localization. Work in this literature often considers only two possibilities: broad distribution or localized hubs. Our data suggest the answer may instead be “both,” which could explain the prevalence of both findings in the literature. One concrete implication of the present work, then, is that the field should more seriously consider such a hybrid distribution as a third possibility. The supplementary beta analyses are consistent with this framing: beta-band RSIs again exhibited a broadly distributed pattern, (and regional concentrations largely mirror those of the high gamma data in sensorimotor regions, where the relationship between beta and high gamma is known to be inverted (*94–96*)).

Another intriguing result is that, despite the near complete lack of overlap between the sentence>list and active≠passive tests at the electrode level, these tests were perfectly aligned at the region level. That is, the sentence>list comparison, which was meant to identify information at the sentence level, was significant in IFG, MFG, MTG, and IPL (Fig. 1D). These are exactly the same regions that were significant for the two sentence-level RSIs: event semantics (MTG & IPL) and structure (IFG & MFG) (see Fig. 4A). Conversely, the two regions that were *not* identified by tests for higher-order language – STG and SMC – were exactly those with above-chance sensitivity to (sub)lexical information. This is precisely the type of alignment we initially anticipated, a prediction which motivated intersecting the two tests. Crucially, however, the ROI-level alignment was driven by mostly distinct sets of electrodes within each region (e.g., different IFG electrodes for sentence>list vs. active≠passive). It remains unclear how and why these tests aligned in ROI analyses without overlap at the electrode level. We suspect that clarity will come from future work that carefully disentangles the many constructs targeted by each of these tests.

### 2.3 Information above and below the word level

Finally, our data reveal a potential distinction in how the brain processes different kinds of information. While (sub)lexical processing was strongly correlated with degree of neural activity, differences associated with sentence structure and event semantics did not show such a relationship. To end on a broader note, we offer a speculative, general hypothesis that could account for this pattern. Specifically, we suggest that this distinction may belie a deeper taxonomic generalization. Consider the fact that (sub)lexical processing is not alone: low-level sensorimotor processes in general show a similar relationship with neural activity, including vision (*126,127*), audition (*128,129*), and motor movement (*130, 131*). We speculate that the divergence between lexical information on one hand and syntactic and event semantic information on the other may be consistent with a broader dichotomy, whereby lower-level systems encode information in a way that elevates activity, while higher-order cognition does not.

Such an account may go some distance to explaining why cognitive neuroscience has progressed more slowly than sensory neuroscience. Just as low-level information shares encoding properties, higher-order processing may in general be encoded in neural populations with varying degrees of overall neural activity. Such a divergence in coding schemes could derive from any number of differences between low- and high-level cognitive systems, including their spatial distribution, evolutionary age, or informational complexity. One possibility is that because higher-level representations are generally sustained over longer periods of time, they require a metabolically efficient neural code. In contrast, lower-level representations like phonemes are relatively short-lived (*132, 133*), and higher firing rates may therefore be achieved without depleting resources. Another possibility relates to the concentration of neural representations of lower- vs. higher-level representations. The neurons that encode a particular pitch in auditory cortex are focally represented in a small patch of cortex. When firing synchronously, this would result in a large local signal (elevated activity, as in the EPE). For a distributed representation, even if encoded by the same number of neurons, if those neurons are spread throughout cortex, then their signal at any given site will be comparatively weak, and the local activity levels will not necessarily show elevation, as we observed. Such an account would be consistent with previous findings of microcircuitry spread out across regions (*36*). It may also help explain the potential spatial pattern among the six electrodes that did show overlap between the sentence>list and active≠passive contrasts, five of which appeared in prefrontal cortex. This region is associated with domain-general executive functions such as working memory, attention, decision-making, and motor planning, which are evolutionarily older processes that exhibit regional localization and may therefore be more likely to scale with activity magnitude, making them more similar in coding properties to (sub)lexical and sensorimotor systems than to higher-order linguistic representations. Under this view, the limited prefrontal overlap highlights a class of computations for which activity magnitude and information content remain coupled. Taken together, our findings provide a new window into the neural underpinnings of sentence production, using high-resolution ECoG data and a controlled production experiment to provide evidence for a hybrid spatial organization and a magnitude-independent coding scheme for higher-order linguistic processing.

## 3 Materials and Methods

### 3.1 Participants

Ten neurosurgical patients undergoing evaluation for refractory epilepsy participated in the experiment (3 female, mean age: 30 years, range: 20 to 45). All ten were implanted with electrocorticographic grids and strips and provided informed consent to participate. All consent was obtained in writing and then requested again orally prior to the beginning of the experiment. Electrode implantation and location were guided solely by clinical requirements. Eight of the participants were implanted with standard clinical electrode grid with 10 mm spaced electrodes (Ad-Tech Medical Instrument, Racine, WI). Two participants consented to a research hybrid grid implant (PMT corporation, Chanassen, MN) that included 64 additional electrodes between the standard clinical contacts (with overall 10 mm spacing and interspersed 5 mm spaced electrodes over select regions), providing denser sampling but with positioning based solely on clinical needs. The study protocol was approved by the NYU Langone Medical Center Committee on Human Research.

### 3.2 Data collection and preprocessing

Participants were tested while resting in their hospital bed in the epilepsy monitoring unit. Stimuli were presented on a laptop screen positioned at a comfortable distance from the participant. Participants’ voice was recorded using a cardioid microphone (Shure MX424). Inaudible TTL pulses marking the onset of a stimulus were generated by the experiment computer, split, and recorded in auxiliary channels of both the clinical Neuroworks Quantum Amplifier (Natus Biomedical, Appleton, WI), which records ECoG, and the audio recorder (Zoom H1 Handy Recorder). The microphone signal was also fed to both the audio recorder and the ECoG amplifier. These redundant recordings were used to sync the speech, experiment, and neural signals.

The standard implanted ECoG arrays contained 64 macro-contacts (2 mm exposed, 10 mm spacing) in an 8×8 grid. Hybrid grids contained 128 electrode channels, including the standard 64 macro-contacts and 64 additional interspersed smaller electrodes (1 mm exposed) between the macro-contacts (providing 10 mm center-to-center spacing between macro-contacts and 5 mm center-to-center spacing between micro/macro contacts, PMT corporation, Chanassen, MN). The FDA-approved hybrid grids were manufactured for research purposes, and a member of the research team explained this to patients during consent. The ECoG arrays were implanted on the left hemisphere for all ten participants. Placement location was solely dictated by clinical care.

ECoG was recorded at 2,048 Hz, which was decimated to 512 Hz prior to processing and analysis. After rejecting electrodes with artifacts (i.e., line noise, poor contact with the cortex, and high amplitude shifts), we subtracted a common average reference (across all valid electrodes and time) from each individual electrode. Electrodes with interictal and epileptiform activity were removed from the analysis. We then extracted the envelope of the high gamma component (the average of three log-spaced frequency bands from 70 to 150 Hz) from the raw signal with the Hilbert transform. Beta activity was quantified as the envelope of the 12 to 30 Hz band.

The signal was separately epoched locked to stimulus (i.e., cartoon images) and production onsets for each trial. The 200 ms silent period preceding stimulus onset (during which patients were not speaking and fixating on a cross in the center of the screen) was used as a baseline, and each epoch for each electrode was normalized to this baseline’s mean and standard deviation (i.e., z-scored).

### 3.3 Experimental Design

#### Procedure

The experiment was performed in one session that lasted approximately 40 minutes and was approved by the NYU Institutional Review Board (IRB). Stimuli were presented in pseudo-random order using PsychoPy (*134*). All stimuli were constructed using the same 6 cartoon characters (chicken, dog, Dracula, Frankenstein, ninja, nurse), chosen to vary along many dimensions (e.g., frequency, phonology, number of syllables, animacy, proper vs. common) to facilitate identification of lexical information at analysis.

The experiment began with two familiarization blocks. In the first block (6 trials), participants saw images of each of the six cartoon characters once with labels (*chicken, dog, Dracula, Frankenstein, ninja, nurse*) written beneath the image. Participants were instructed to read the labels aloud, after which the experimenter pressed a button to go to the next trial. In the second block, participants saw the same six characters one at a time, twice each in pseudo-random order (12 trials), but without labels. They were instructed to name the characters out loud. After naming the character, the experimenter pressed a button revealing the target name to ensure learning of the correct labels. Participants then completed the picture naming block (96 trials). As before, characters were presented in the center of the screen, one at a time, but no labels were provided.

Next, participants performed a sentence production block (60 trials), starting with two practice trials. Participants were instructed that there were no right or wrong answers, we want to know what their brain does when they speak naturally. On each trial, participants saw a 1 s fixation cross followed by a written question, which they were instructed to read aloud to ensure attention. After another 1 s fixation cross, a static cartoon vignette appeared in the center of the screen depicting two of the six characters engaged in a transitive event. Participants were instructed to respond to the question by describing the vignette. The image remained on the screen until the participant completed the sentence, at which point the experimenter pressed a button to proceed. After the first 12 trials, the target sentence (i.e., an active sentence after an active question or passive sentence after a passive question) appeared in text on the screen and participants were instructed to read aloud “the sentence we expected [them] to say” to implicitly reinforce the link between the syntax of the question and the target response. If participants appeared to interpret these as corrections, the experimenter reminded them that there were no right or wrong answers.

We interleaved two picture naming trials between each sentence production trial in order to reduce task difficulty and facilitate fluent sentence production. The picture naming trials involved the two characters that would be engaged in the subsequent vignette, presented in a counterbalanced order such that on half of trials they would appear in the same order as in the expected sentence response, and in the opposite order on the other half.

Next, participants performed the listing block. List production trials were designed to parallel sentence production trials. Each trial began with a 1 s fixation cross and then an arrow pointing either left or right appeared for 1 s in the center of the screen. After another 1 s fixation cross, a cartoon vignette (the exact same stimuli) appeared on the screen. Participants named the two characters in the vignette from left to right or from right to left, according to the direction of the preceding arrow. As in sentence production trials, each list production trial was preceded by two picture naming trials with the two characters that would appear in the subsequent vignette, again in counterbalanced order.

Between each block, participants were offered the opportunity to stop the experiment if they did not wish to continue. One participant stopped before the list production block, providing only picture naming and sentence production data. The other nine participants completed all three blocks. These nine were offered the opportunity to complete another picture naming block and another sentence production block. Six consented to another picture naming block and two consented to another sentence production block.

#### Stimulus Design, Randomization, and Counterbalancing

Picture naming stimuli consisted of images of the 6 characters in pseudo-random order so that each consecutive set of 6 trials contained all 6 characters in random order. This ensured a relatively even distribution of characters across the block, and that no character appeared more than twice in a row. Characters were pseudorandomly depicted facing 8 orientations: facing forward, backward, left, right, and at the 45^◦^ angle between each of these.

Sentence production stimuli consisted of a written question and a static cartoon vignette. Questions were manipulated so half were constructed with passive syntax and half with active. All questions followed the scheme: “Who is [verb]-ing whom?” or “Who is being [verb]-ed by whom?”. There were 10 verbs: *burn, hit, hypnotize, measure, poke, scare, scrub, spray, tickle, trip*. Each verb was used to create 3 vignettes involving 3 characters, counterbalanced so each character was the agent (i.e., active subject) in one vignette and the non-agent (i.e., active object) in one vignette. Each of these three vignettes was shown twice in the sentence production block, once preceded by an active question and once by a passive question to prime active and passive responses, respectively (*88*). Vignettes were flipped around the vertical axis the second time they appeared so that the character that was on the left in the first appearance was on the right in the second appearance and vice versa. This was counterbalanced so that on half of the trials in each structure condition (active/passive) the subject was on the left. List production stimuli similarly consisted of the same 60 vignettes, also presented in pseudorandom order and counterbalanced across conditions (i.e., arrow direction).

### 3.4 Data Coding and Inclusion

Speech was manually transcribed and word onset times were manually recorded using Audacity (*135*) to visualize the audio waveform and spectrogram. Picture naming trials were excluded if the first word uttered was not the target word (e.g., “Frankenstein – I mean Dracula”). Sentence trials were excluded if the first word was incorrect (i.e., “Frankenstein” instead of “Dracula,” regardless of syntactic structure) or if the meaning of the sentence did not match the meaning of the depicted scene; no sentences were excluded because the structure did not match that of the prime (i.e., question). Sentences were coded as active or passive depending on the structure the patient used, not the prime structure. Listing trials were excluded if the first word was incorrect or if the order did not match that indicated by the arrow.

All patients were included in Figure 1 analyses, however three patients who produced 6 or fewer passive sentences during the sentence production block were excluded prior to any subsequent analyses that involved an active≠passive test (including the RSA and NMF analyses).

### 3.5 Electrode Localization

Electrode localization in subject space, as well as MNI space, was based on coregistering a pre-operative (no electrodes) and postoperative (with electrodes) structural MRI (in some cases, a postoperative CT was employed depending on clinical requirements) using a rigid-body transformation. Electrodes were then projected to the surface of the cortex (preoperative segmented surface) to correct for edema-induced shifts following previous procedures (*136*) (registration to MNI space was based on a nonlinear DARTEL algorithm). Based on the subject’s preoperative MRI, the automated FreeSurfer segmentation (Destrieux) was used for labeling within-subject anatomical locations of electrodes. Plotting to a standard brain was performed using the Mithra toolbox (*137*).

### 3.6 Significance testing and corrections for multiple comparisons in time series data

Statistical tests on time series data were performed independently at each time sample, resulting in the same number of *p*-values as there are samples in the time series. All time series data were sampled at 512 Hz (i.e., a step size of 2 ms). To correct for multiple comparisons we follow (*138, 139*) and establish a conservative criterion for significance for all time series comparisons: an uncorrected *p*-value that remains below .05 for at least 100 consecutive milliseconds. Note that this only applies to time series data; for aggregate measures, like activity within a particular time bin as in Fig. 1C, we used traditional tests and corrections for multiple comparisons (e.g., FDR).

### 3.7 Permutation tests for ROIs

To determine whether activity was significantly above chance for a given ROI (Figs. 1D and 4A), we randomized electrodes’ ROI labels and re-calculated ROI means 1000 times. We derived *p*-values by determining what proportion of these chance means were above the real mean value at each time sample. If the *p*-value was less than .05 for at least 100 consecutive milliseconds (see Sec. 3.6), it was considered significant, denoted by a bar at the bottom of the plot.

### 3.8 Temporal warping

Our analyses focused on the planning period – the time between stimulus and speech onsets, when hierarchical syntactic structure is planned (*29, 81, 84, 93*). The duration of the planning period varied considerably both across and within patients, meaning that cognitive processes become less temporally aligned across trials the farther one moves from stimulus onset in stimulus-locked epochs or speech onset in speech-locked epochs. This was potentially problematic for comparing syntactic structures, as passive trials took longer to plan (median RT: 1,424 ms) than active trials (median RT: 1,165 ms; *p* < 0.001, Wilcoxon rank sum test). The farther from the time lock, the more misaligned active and passive trials would be, and the more likely significant differences would be to reflect temporal misalignment rather than structure-related differences.

Temporal warping reduces such misalignments (*98–100, 140, 141*). Following (*98*), we linearly interpolated the data in the middle of the planning period (from 150 ms post-stimulus to 150 ms pre-speech) for each trial to set all trials’ planning periods to the same duration (Supplementary Fig. S1): the global median per task (1141 ms for sentences; 801 ms for lists; 758 ms for words). Specifically, for each task we started by excluding trials with outlier response times, which we defined as those in the bottom 2.5% or top 5% per participant. We then calculated the median response time per task across participants (1,142 ms for sentences, 800 ms for lists, and 758 ms for words), and for each electrode and each trial, concatenated (a) the stimulus-locked data from 150 ms post-stimulus to 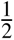 the median reaction time with (b) the production-locked data from 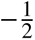 median reaction time to 150 ms pre-speech. We then linearly interpolated this time series to the number of time samples corresponding to the median reaction time minus 300 ms (i.e., the two 150 ms periods following stimulus onset and preceding speech onset). Finally, we concatenated (a) the unwarped data leading up to 150 ms post-stimulus, (b) the warped data from the previous step, and (c) the unwarped data starting 150 ms before speech onset to form the final epochs used in the analyses in Figs. 2, 3, and 4.

Improved temporal alignment leads to better signal-to-noise ratio, which can be seen in higher trial means (*100*). We leveraged this fact as a diagnostic for whether temporal warping in fact improved alignment in our data. For electrodes whose mean high gamma peaked in the warped period (between 150 ms post-stimulus and 150 ms pre-speech), we compared the peak mean values in the unwarped and the warped data. Peak values were significantly higher for the warped data than the unwarped data (Supplementary Fig. S3) for both the sentence and list production tasks (*p* = 0.003 and *p*.008 respectively, Wilcoxon signed rank tests), indicating that warping significantly improved temporal alignment in the data. (There was no significant difference between peak values in the picture naming data (*p* = 0.592). This likely reflects the fact that these trials were relatively short, with a median reaction time of just 758 ms, meaning that warping made relatively minor changes to a smaller number of time samples.)

For comparison, we reproduced Fig. 1 in Supplementary Fig. S2 using the warped data. The results of the statistical tests in these figures were qualitatively identical: The spatial distributions of significant electrodes were nearly identical over time (Fig. 1C and Supplementary Fig. S2C) and the same ROIs (IFG, MFG, MTG, and IPL) showed significantly higher activity for sentences than lists during the planning period (Fig. 1D and Supplementary Fig. S2D).

### 3.9 Representational Similarity Analysis (RSA) and Representational Similarity Indices (RSIs)

To uniquely identify variance associated with event-semantics, (sub)lexical processing, and the active/passive distinction, we used multiple linear regression to model the neural activity from each electrode and at each time sample as a linear combination of event-semantic, (sub)lexical, and structural properties of the sentence the patient was planning/producing. Multiple regression is ideal in this context because it fits coefficients (slopes/betas) that can be used to derive *t*-values and corresponding *p*-values which reflect the *unique* contribution of each independent variable, effectively partitioning variance that can be unambiguously ascribed to just one term (*142*).

However, event semantic and (sub)lexical representations are highly complex and multidimensional, requiring a choice of which dimensions/features to use in a multiple regression model. To avoid this scenario, we leveraged a Representational Similarity Analysis-style approach and modeled pairwise differences between trials (*101–104*). This resulted in just one vector per construct representing pairwise trial differences in sentence structure, event semantics, and word content.

#### Linguistic and Neural Dissimilarity Models

The structure term was binary: 1 if one trial was active and the other was passive and 0 otherwise. We modeled event-semantic representations using outputs from GPT-2’s 8th hidden layer, where embeddings correlate most highly with neural activity (*143*). Note that these embeddings should not be interpreted as isolating “pure” event semantics in a theoretical sense, but rather as providing a high-dimensional representation of sentence-level meaning. Specifically, for each unique stimulus vignette, we fed the corresponding sentence with active syntax into GPT-2 and averaged the embeddings in layer 8 for all of the words in the sentence. By inputting only active sentences (rather than a combination of actives and passives depending on the structure of the specific trials), we ensured that the resulting embedding-based dissimilarity model was orthogonal to the active/passive manipulation, such that variance captured by this term reflects differences in sentence-level meaning rather than syntactic structure or surface word order.

To quantify event-semantic dissimilarity between trials, we correlated the vectors corresponding to each pair of trials’ stimuli. Prior to modeling, correlation coefficients (*r*) were centered, scaled, and multiplied by −1 so that a value of 1 corresponded to more dissimilar meanings, and −1 meant that the two trials had the same exact stimulus. (Each stimulus appeared twice in each sentence production block, once with an active prime and once with a passive prime.)

The (sub)lexical term was also binary: 1 if the first word of the two trials’ sentences were different, 0 if they were the same. This encoding scheme was meant to absorb variance associated with a host of extraneous linguistic features at the lexical and sub-lexical levels including phonetics, phonology, articulatory/motor information, auditory feedback, lemma-level representations, and lexical semantics (the meaning of individual words rather than the global event meaning).

We also included differences in response time (RT) as a covariate in the model, as this is known to index other extraneous non-linguistic features like difficulty that might be correlated with our structural manipulation. This term was quantified as the absolute value of the difference in the log of each trial’s reaction times, so that higher values corresponded to bigger differences in reaction times.

We then modeled pairwise trial differences in neural activity (the magnitude of the difference between *z*-scored high gamma activity at a given sample for a given electrode) as a linear combination of these four dissimilarity (D) terms:

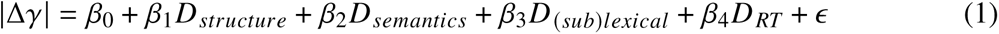

#### Electrode Significance

The data used in the RSA regression did not clearly meet the assumptions of standard linear regressions: it was not clear that the residuals should be normally distributed (high gamma activity is gamma-distributed), and imbalances in the datasets due to trial exclusions and the varying effectiveness of priming across patients were exacerbated by the implementation of pairwise trial differences. To err on the side of caution, rather than directly interpreting model statistics, we ran a permutation analysis, shuffling the neural activity with respect to linguistic features in the original datasets, reconstructing the neural dissimilarity models, and re-running the linear regression at each time sample 1000 times.

To assess whether a given electrode was significant for the structure, semantics, and (sub)lexical terms, we started by calculating the *t*-values corresponding to the each term’s coefficient, which corresponds to the amount of evidence against the null hypothesis. Then we smoothed the real and shuffled *t*-values over time with a 100 ms boxcar function. Finally, we *z*-scored the real *t*-values by time sample and electrode with respect to the 1000 *t*-values from shuffled models in the same time sample and electrode. The resulting *z*-values reflect an estimate of evidence against the null that is standardized across patients and electrodes and independent of the number of trials. Significant electrodes for each representation were defined as those which maintained a *z*-value of at least 1.645 (corresponding to a 1-tailed *p*-value of less than .05) for at least 100 consecutive milliseconds.

#### Representational Similarity Indices (RSIs)

To derive RSIs, we performed additional transformations on these *z*-values to make them more interpretable and conservative. First, We scaled *z*-scores, dividing by 2.326, which corresponds to a *p*-value of .01. Consequently, values greater than 1 could be interpreted as very likely to reflect positive results, and values less than 1 were likely to reflect negative results. Next, to reduce the possibility of extremely high values having an undue influence on aggregate statistics and NMF, we applied a logarithmic penalty to values greater than 1, replacing them with *log*(*x*) + 1. Similarly, to reduce values more likely to correspond to negative results, we shrunk low values toward 0 by replacing values less than 1 with (*e*^*x*^), and then cubing the resulting values (i.e., (*e*^*x*^)^3^) to impose an even more severe penalty on low values. Notably, this renders all values non-negative, naturally resolving issues related to the uninterpretability of negative relationships in RSA (*101, 144*) and facilitating the use of NMF for subsequent clustering. Finally, we re-scaled values by multiplying them by 2.326 (undoing Step 1) so that the final RSI scale more closely matched the *z* scale in interpretation. Like *z*-scores, RSI values over 2.326 can be safely interpreted as significant at the *α* = 0.01 threshold.

### 3.10 Non-negative Matrix Factorization (NMF)

We used NMF to cluster electrodes according to prototypical patterns in the data (*145, 146*). In the first clustering analysis (Fig. 4), we analyzed RSIs and mean neural activity from all three tasks for all electrodes that were either “active” (non-zero neural activity; *p* < 0.05 for 100 ms, Wilcoxon signed rank test) or inactive, but with at least one significant RSI (*p* < 0.05 for 100 ms, permutation test).

High gamma activity was in units of standard deviations (i.e., *z*-scores), while RSIs were *z*-scores that had been smoothed and transformed to be non-negative, with extreme values in both directions shrunk (see Section 3.9). To ensure that NMF weighted the two types of information the same way, we applied the same transformation used to create the RSIs (Sec. 3.9) to the neural data prior to performing NMF.

We then concatenated these three time series (i.e., mean high gamma per task) and the three RSI time series (i.e., structure, event semantics, and (sub)lexical), excluding pre-stimulus time samples and those after 500 ms post-speech onset. This resulted in a matrix of dimensionality 741 (electrodes) × 4364 (time samples from 3 high gamma and 3 RSI time series).

We fed this matrix to the nmf() function in R (*NMF* library v0.25 (*147*)) using the Brunet algorithm (*148*) and 50 runs to mitigate the risk of converging on local minima. We tried a range of model ranks: 3 to 9. The optimal rank (5) was determined by finding the elbow in the scree plots.

## Acknowledgments

We would like to thank Leyao Yu, Amirhossein Khalilian-Gourtani, and our anonymous reviewers for invaluable input on this project.

## Author contributions

Each author’s contributions are listed below using the CRediT model.

Conceptualization: AMM, AF

Data curation: AMM, OD, WKD, PD, DF

Formal analysis: AMM

Funding Acquisition: AMM, AF

Investigation: AMM

Methodology: AMM, AF

Project administration: AMM, AF

Supervision: AF

Software: AMM, AF

Validation: AMM

Visualization: AMM

Writing – original draft: AMM

Writing – review & editing: AMM, AF

## Data, code, and materials availability

The data set generated during the current study will be made available from the authors upon request and documentation is provided that the data will be strictly used for research purposes and will comply with the terms of our study IRB. The code is available on OSF at https://doi.org/10.17605/OSF.IO/6QCWK. Materials used during this study can be made available upon request via the corresponding author.

This study was approved by the NYU Grossman School of Medicine Institutional Review Board (IRB; approved protocol s14-02101) which operates under NYU Langone Health Human Research Protections. The research study was performed in accordance with the Department of Health and Human Services policies and regulations at 45 CFR 46. NYU Langone Health uses specific Federalwide Assurance (FWA) numbers for human research protections; the main NYU Grossman School of Medicine FWA is 00004952. Before obtaining consent, a clinical staff member confirmed that all participants had cognitive capacity to provide informed consent. Participants provided informed consent orally and in writing before beginning study procedures. They were informed that their participation was strictly voluntary, and would not impact clinical care. Participants were further informed that they were free to withdraw participation in the study at any time. All procedures for this study were conducted in accordance with the Declaration of Helsinki.

## Supplementary Text

### Data Warping

Supplementary Figures S1, S2, and S3 show the methods and efficacy checks for our temporal warping procedure (see Sec. 3.8). Supplementary Figure S3 analyses were performed on the maximum values of electrodes’ trial means. Prior to finding the maximum, we took the absolute value of the trial mean to capture electrodes with negative peaks (reflecting suppression) that might have been enhanced by warping. We included only those electrodes that peaked during the warped period (between 150 ms post-stimulus and 150 ms pre-speech), as other electrode’s maxima remained unchanged.

**Figure S1:**
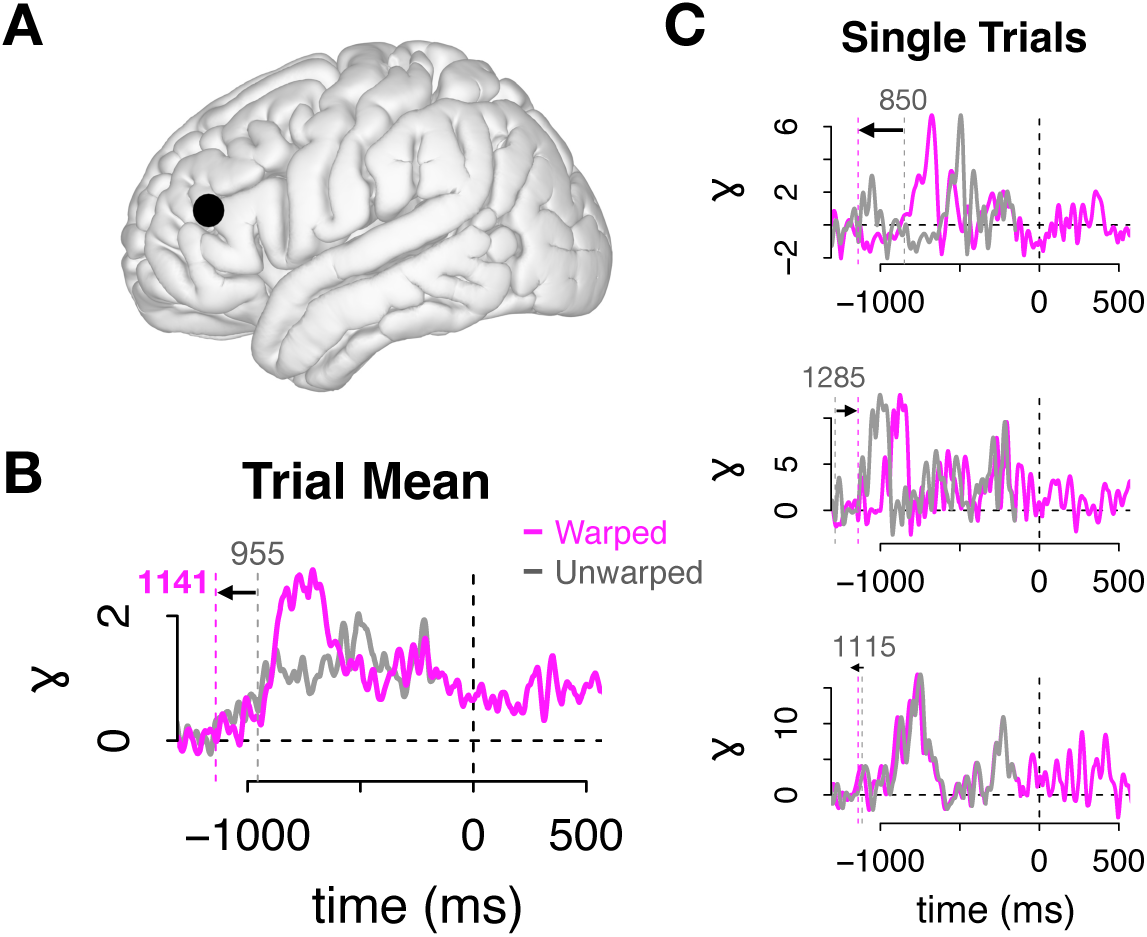
Example of warped versus unwarped high gamma activity in sentence production for a single electrode. Warped and unwarped sentence production data from a sample electrode. The data in each trial between 150 ms post stimulus and 150 ms pre-speech were linearly interpolated to set the duration of the planning period to the global median per task (1142 ms for sentence production) (*98*). (A) Sample electrode localization in MFG. (B) The mean of this electrode’s warped (pink) and unwarped (grey) trials. Prior to warping, this patient’s median sentence response time was 995 ms; after warping it was 1,141 ms: the median sentence production response time across patients. The peak of the warped data was higher than the unwarped peak, a sign that warping resulted in better temporal alignment and consequently higher signal-to-noise ratio. (C) Three sample trials: warped (pink) and unwarped (grey) data.

**Figure S2:**
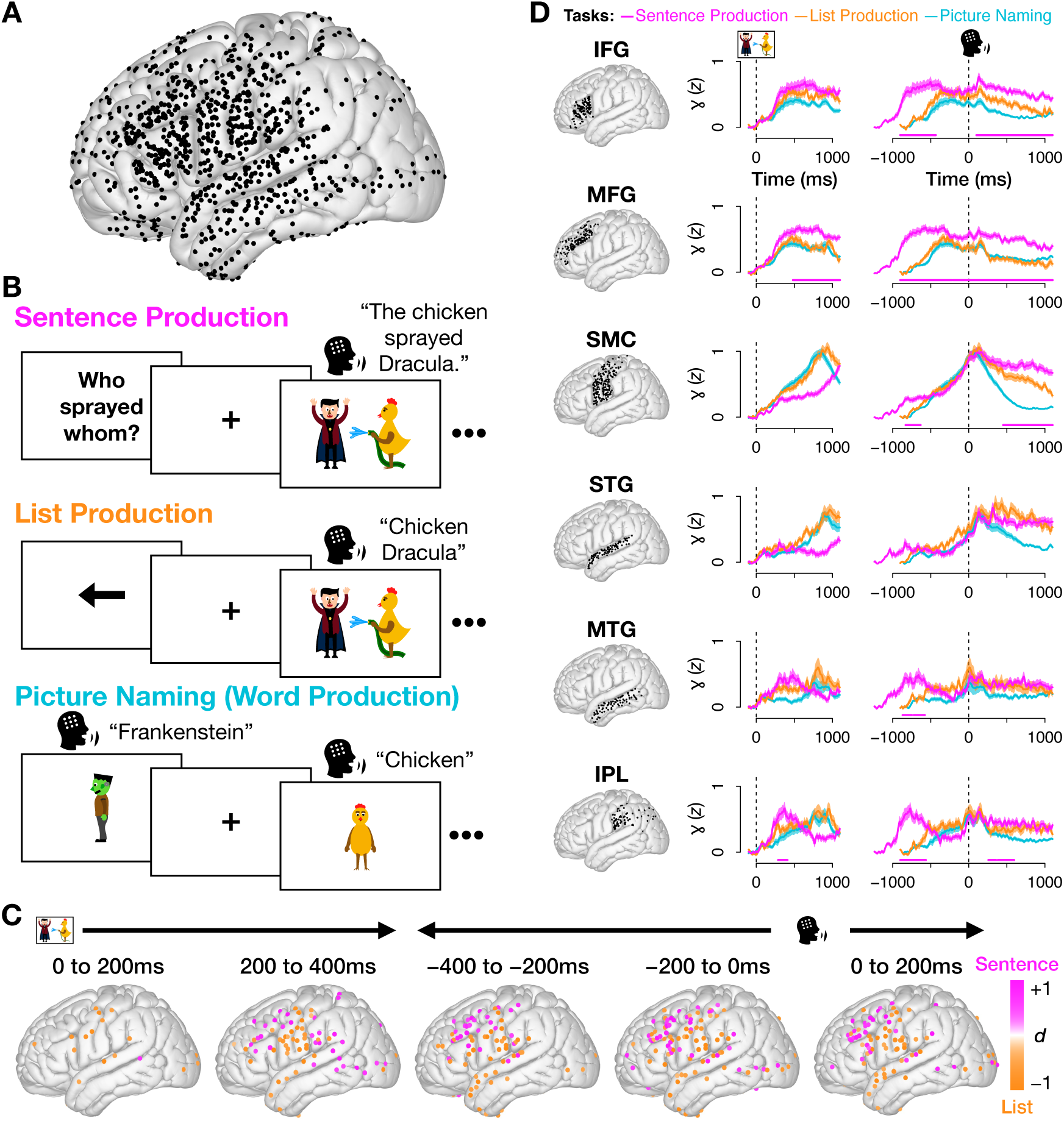
Sentence–list high gamma effects are robust to temporal warping of the planning period. Here we replicate Figure 1 using warped data. (A) Our coverage (unchanged). (B) Experimental paradigm (unchanged). (N.B.: The cartoon figures here and throughout are not the exact images used in the experimental stimuli, but are visually similar. They were created by the authors (*149*) for publication.) Results are qualitatively identical: (C) the spatial distribution of electrodes with significant differences between sentences and lists over time was nearly identical and (D) both identify IFG, MFG, MTG, and IPL as having significantly higher sentence than list activity during the planning period.

**Figure S3:**
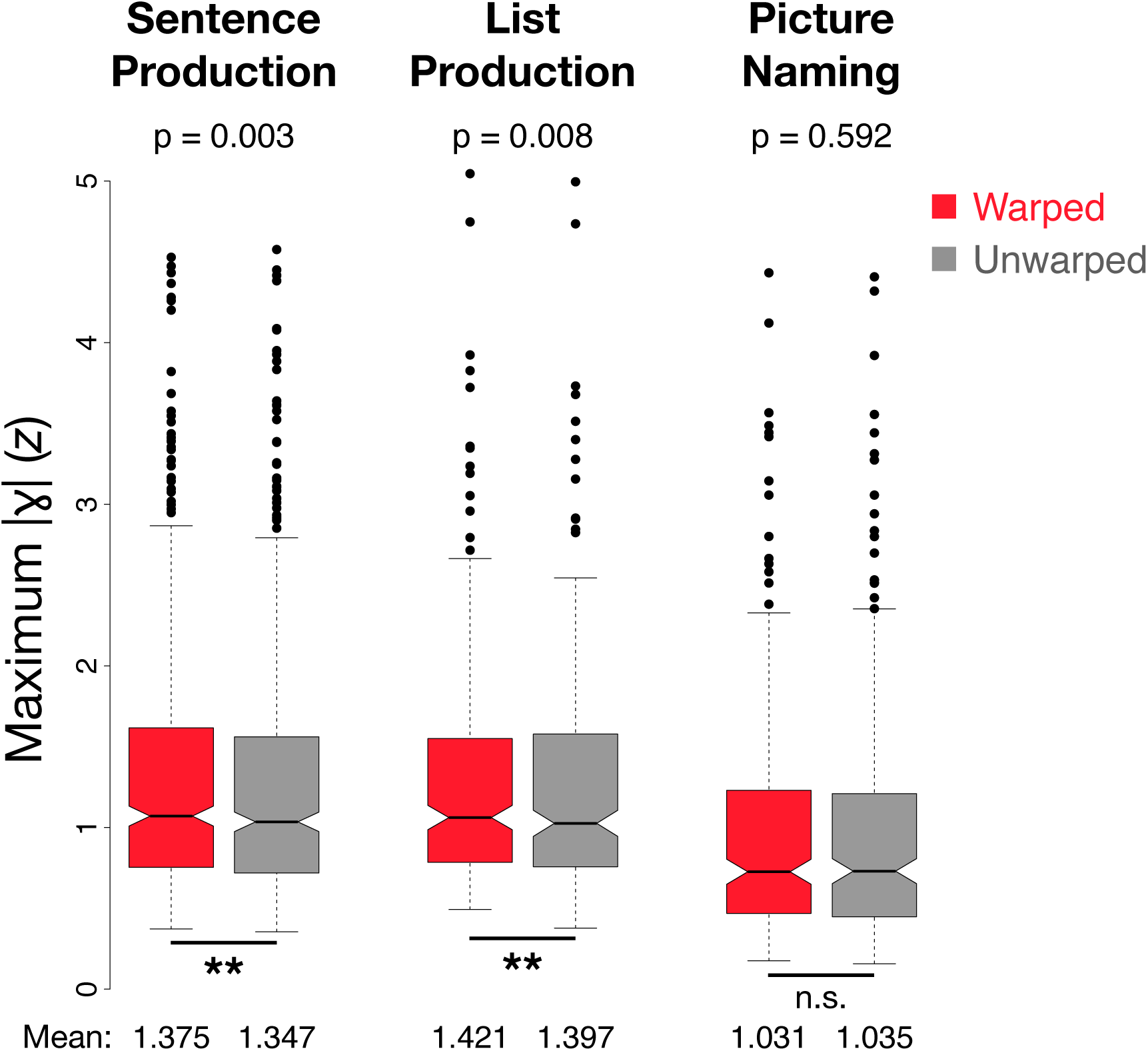
Temporal warping improves alignment of high gamma during sentence and list planning. Peak high gamma activity for each electrode (dot) in the warped (red) and unwarped (grey) data, by task. Warping resulted in significantly higher peaks for sentence production (*p* = 0.003, Wilcoxon signed rank test) and list production (*p* = 0.008, Wilcoxon signed rank test), evidence that it successfully improved the temporal alignment of trials (*100*). No effect was found for picture naming (*p* = 0.592).

### Analyses of Beta Activity

Supplementary Fig. S4 replicates the Figure 1 analyses but using beta activity (12-30 Hz). Beta is another important frequency band in cognition (*91, 92*). In Supplementary Fig. S5, we test the hypothesis that beta activity tracks structural differences more clearly than high gamma. We present three replications of the analysis in Fig. 3A (copied in Supplementary Fig. S5A). Analyses appear in a 2 × 2 factorial design: regressing either beta or high gamma activity over RSIs, where RSIs were derived from RSA performed on either high gamma (left) or beta (right) activity. Across all four sets of results, the only significant finding is that reported in the main text: a positive relationship between mean sentence high gamma and (sub)lexical signal in the high gamma trial activity (*p* < 0.001, linear regression).

To determine whether representational encoding in beta activity might be anatomically localized more than in high gamma, we replicated the analyses in Fig. 2 using beta. We analyzed the resulting “beta RSIs” by region of interest (Supplementary Fig. S6A). There was significant evidence for sensitivity to structure in SMC and, as in the high gamma data, IFG (*p* < 0.05 for 100 ms, permutation test), but unlike high gamma there was no evidence that beta activity was sensitive to the active/passive contrast in MFG. We also plotted electrodes with significant beta RSIs on cortex (Supplementary Fig. S6B), revealing distributed networks similar to those observed in high gamma.

**Figure S4:**
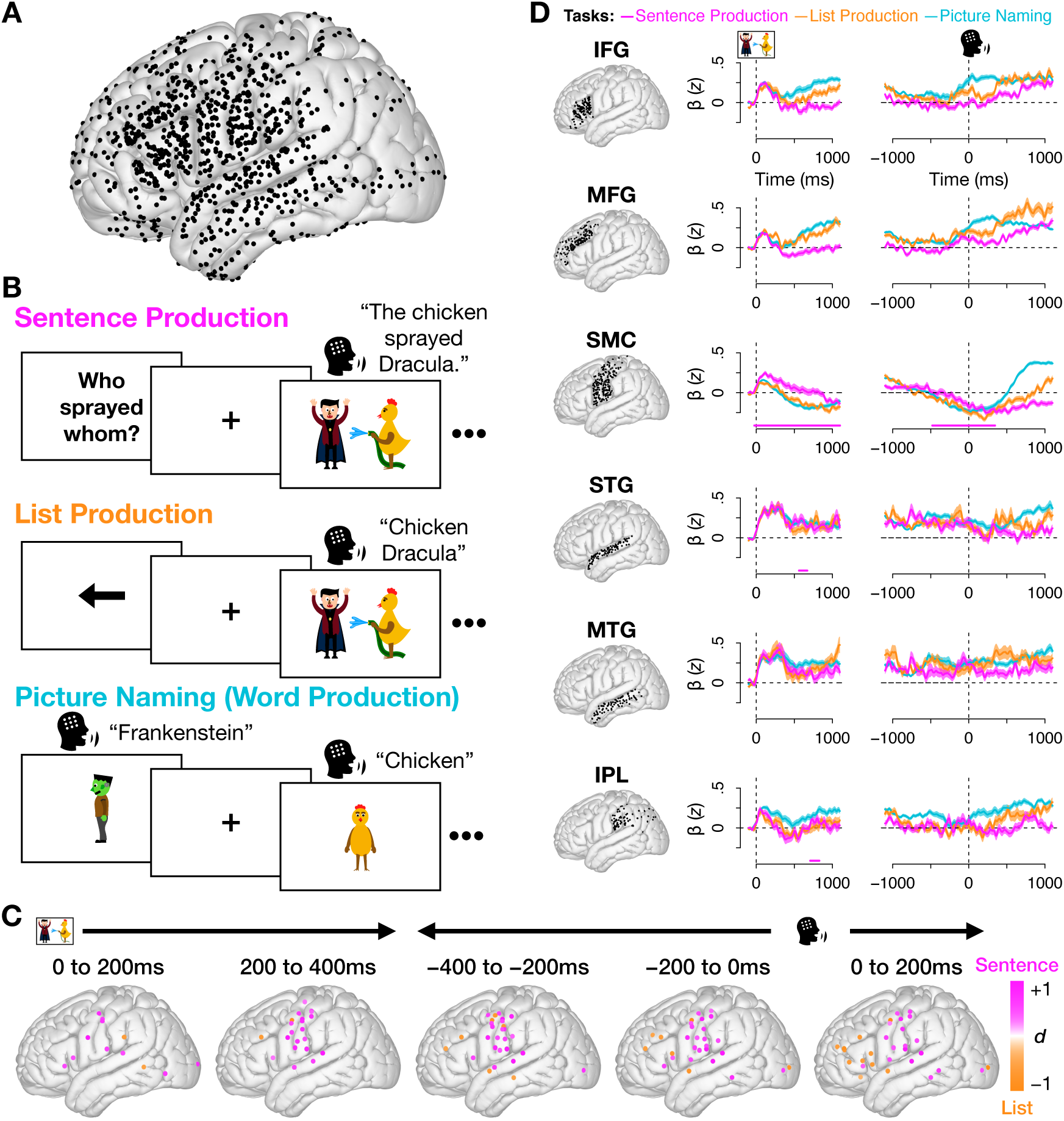
Beta-band activity shows suppression corresponding to high gamma increases. Replication of Figure 1 using beta activity (12-30 Hz; unwarped data). (C) The distribution of electrodes that are significantly greater for sentences than lists was largely reversed from the high gamma activity. This likely reflects a well-documented phenomenon where activity in high gamma is often coupled with beta suppression, particularly in sensorimotor regions, leading to effects in the reverse direction in beta (*150*, *151*). (D) SMC, STG, and IPL show significantly higher beta activity for sentences than lists, possibly also reflecting beta suppression corresponding to increases in high gamma during speech and auditory feedback.

**Figure S5:**
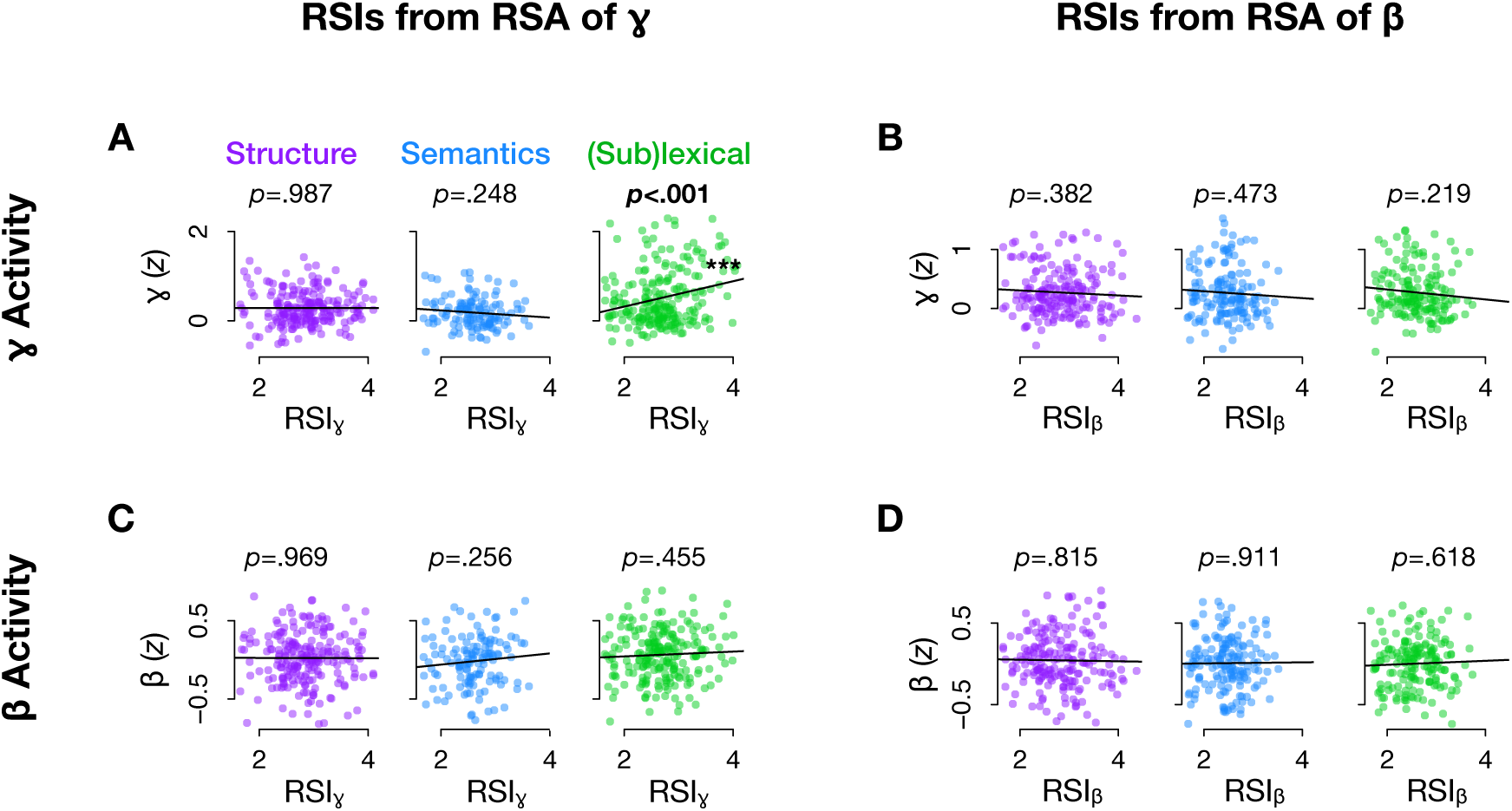
Across beta and high gamma, mean activity tracks (sub)lexical but not higher-order linguistic processing. Four attempts at uncovering a relationship between mean neural activity during sentence production and linguistic processes. Only one significant relationship was found across all analyses: mean high gamma activity was positively related to high gamma (sub)lexical processing (*p* < 0.001). (A) Mean high gamma activity vs. linguistic RSIs encoded in high gamma trial activity (panel is identical to Fig. 3A). (B) Mean high gamma activity vs. linguistic RSIs encoded in beta trial activity. (C) Mean beta activity vs. linguistic RSIs encoded in high gamma trial activity. (D) Mean beta activity vs. linguistic RSIs encoded in beta trial activity.

**Figure S6:**
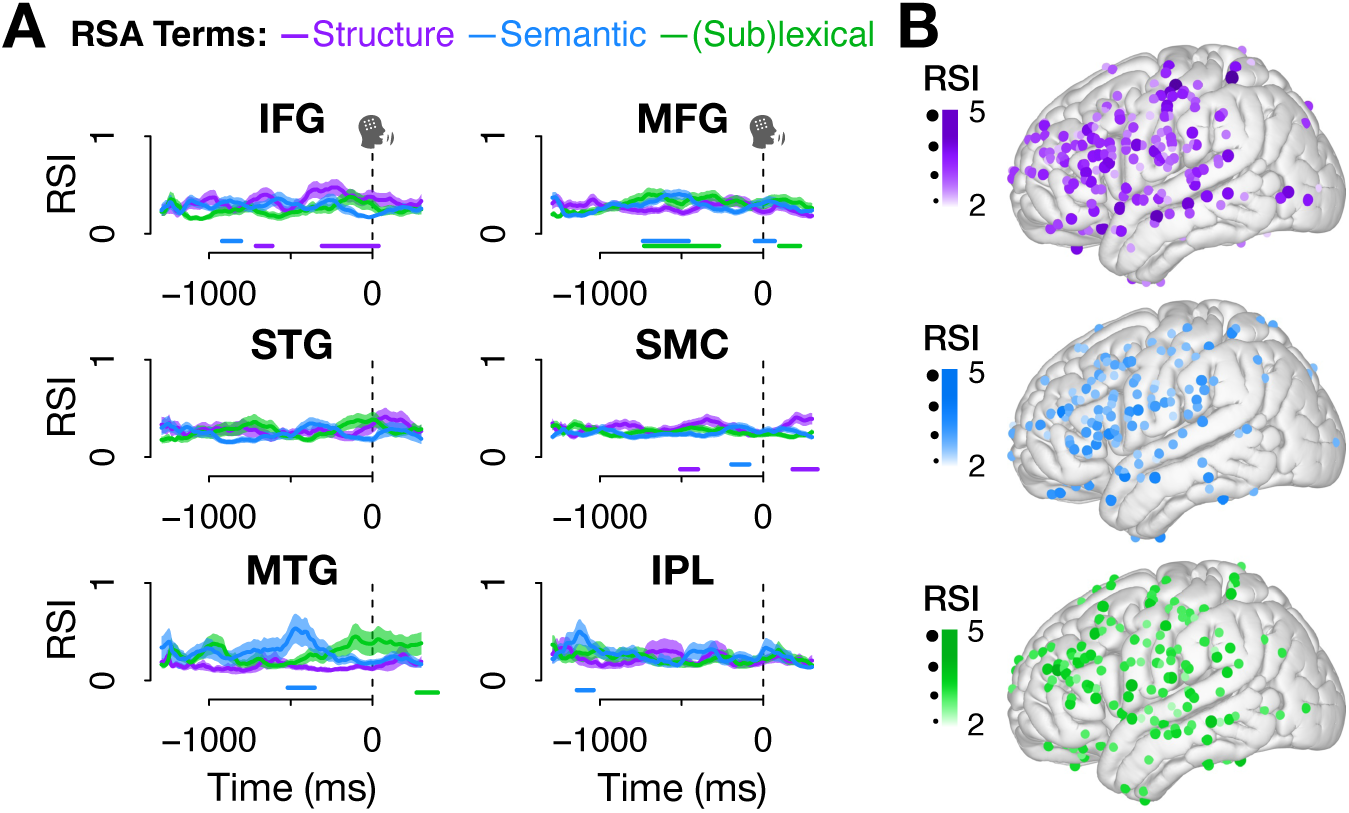
Distributed beta-band sensitivity to linguistic information parallels high gamma RSIs. Distribution of RSIs calculated from RSA on beta activity (see parallels for high gamma in Fig. 2). (A) Mean and standard error by region shows significant structure effects in IFG and SMC (*p* < 0.05 for 100 ms, permutation test), but there was no evidence for sensitivity to structure in MFG as there was in the high gamma activity. (B) Significant electrodes (*p* < 0.05 for 100 ms; one-tailed permutation test) per RSI again show a broadly distributed pattern for all three RSIs, as in high gamma.

### Anatomy of RSIs and their overlap

Figure S7 shows the distribution of electrodes which were significant for each RSI (and each combination of RSIs) during the planning period.

**Figure S7:**
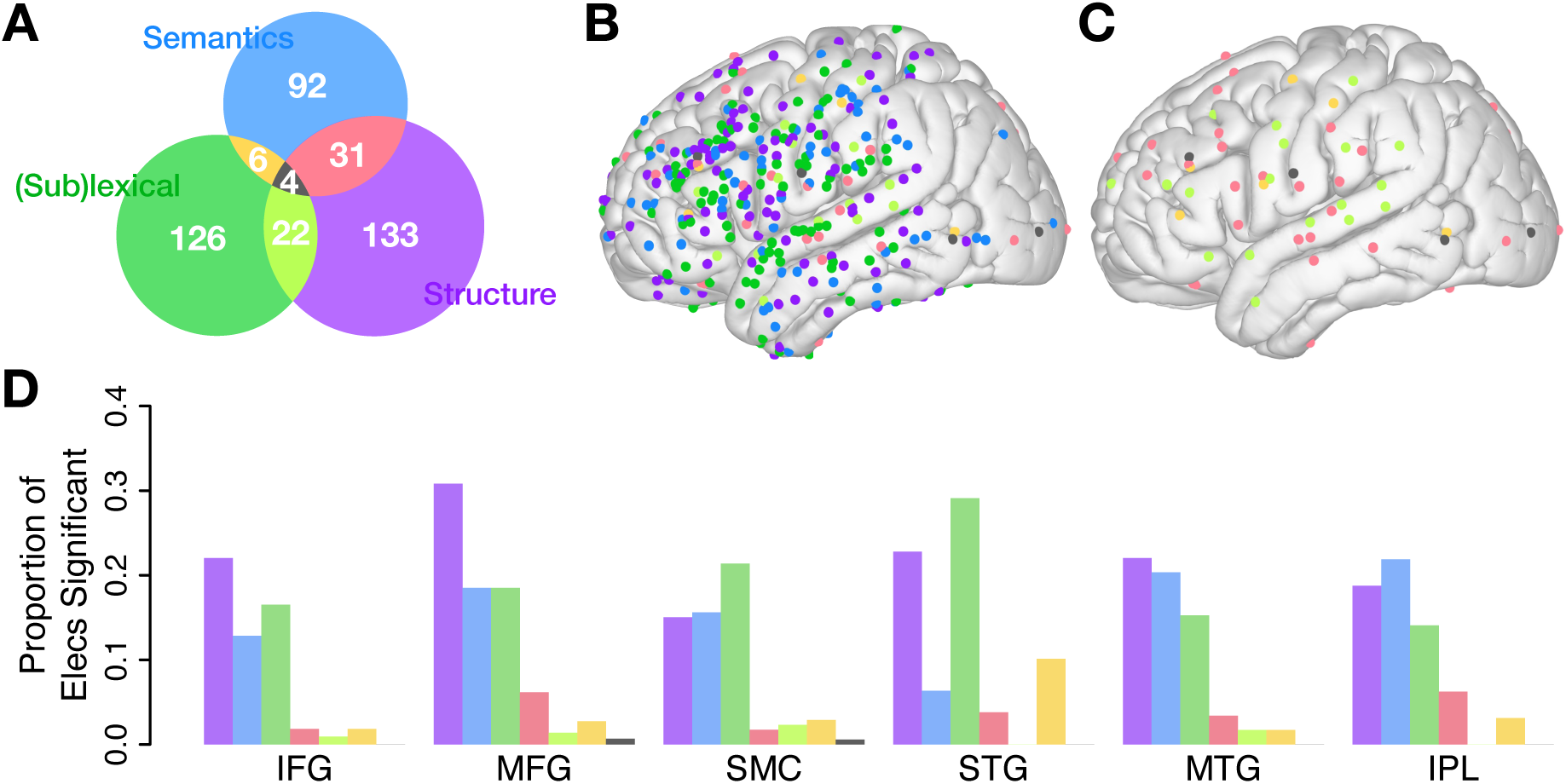
Anatomical distributions of RSIs. (A) The number of electrodes significant for each RSI during the planning period (a replication of Fig. 2E to serve as a color key for the remaining panels in this figure). (B) All electrodes significant for one or more RSI. Colors correspond to the Venn diagram in Panel A, indicating which RSI(s) each electrode was significant for. (C) The distribution of overlapping structural, event semantic, and/or (sub)lexical information (i.e., just electrodes significant for two or three RSIs). (See Fig. 4A-B for the distribution of individual RSIs.) (D) The proportion of analyzed electrodes in each ROI which were significant for each RSI (and combinations therein).

