## Supplemental Materials for "Hybrid spatial organization and magnitude-independent neural coding of linguistic information during sentence production"

**This PDF file includes:**

Supplementary Text
Figures S1 to S7

### Supplementary Text

#### Data Warping

Supplementary Figures S1, S2, and S3 show the methods and efficacy checks for our temporal warping procedure (see Sec. 3.8). Supplementary Figure S3 analyses were performed on the maximum values of electrodes' trial means. Prior to finding the maximum, we took the absolute value of the trial mean to capture electrodes with negative peaks (reflecting suppression) that might have been enhanced by warping. We included only those electrodes that peaked during the warped period (between 150 ms post-stimulus and 150 ms pre-speech), as other electrode's maxima remained unchanged.

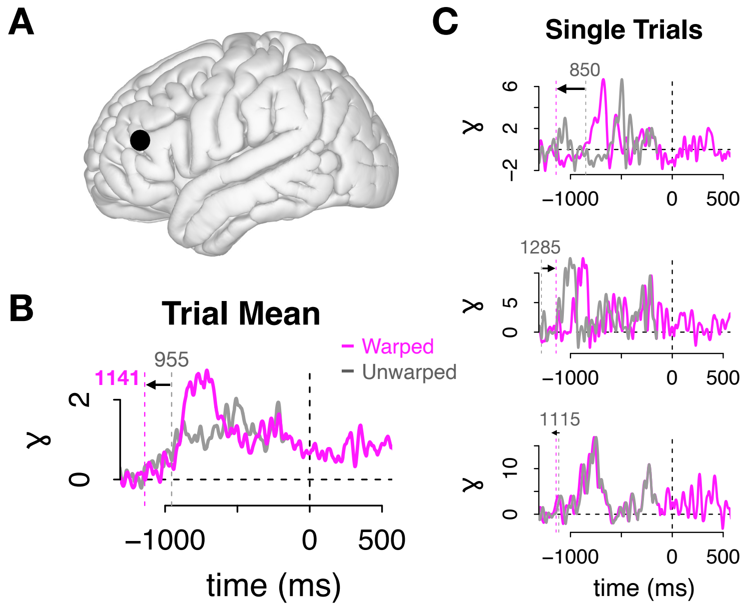


**Figure S1**: **Example of warped versus unwarped high gamma activity in sentence production for a single electrode.** Warped and unwarped sentence production data from a sample electrode. The data in each trial between 150 ms post stimulus and 150 ms pre-speech were linearly interpolated to set the duration of the planning period to the global median per task (1142 ms for sentence production) (*98*). (A) Sample electrode localization in MFG. (B) The mean of this electrode's warped (pink) and unwarped (grey) trials. Prior to warping, this patient's median sentence response time was 995 ms; after warping it was 1,141 ms: the median sentence production response time across patients. The peak of the warped data was higher than the unwarped peak, a sign that warping resulted in better temporal alignment and consequently higher signal-to-noise ratio. (C) Three sample trials: warped (pink) and unwarped (grey) data.

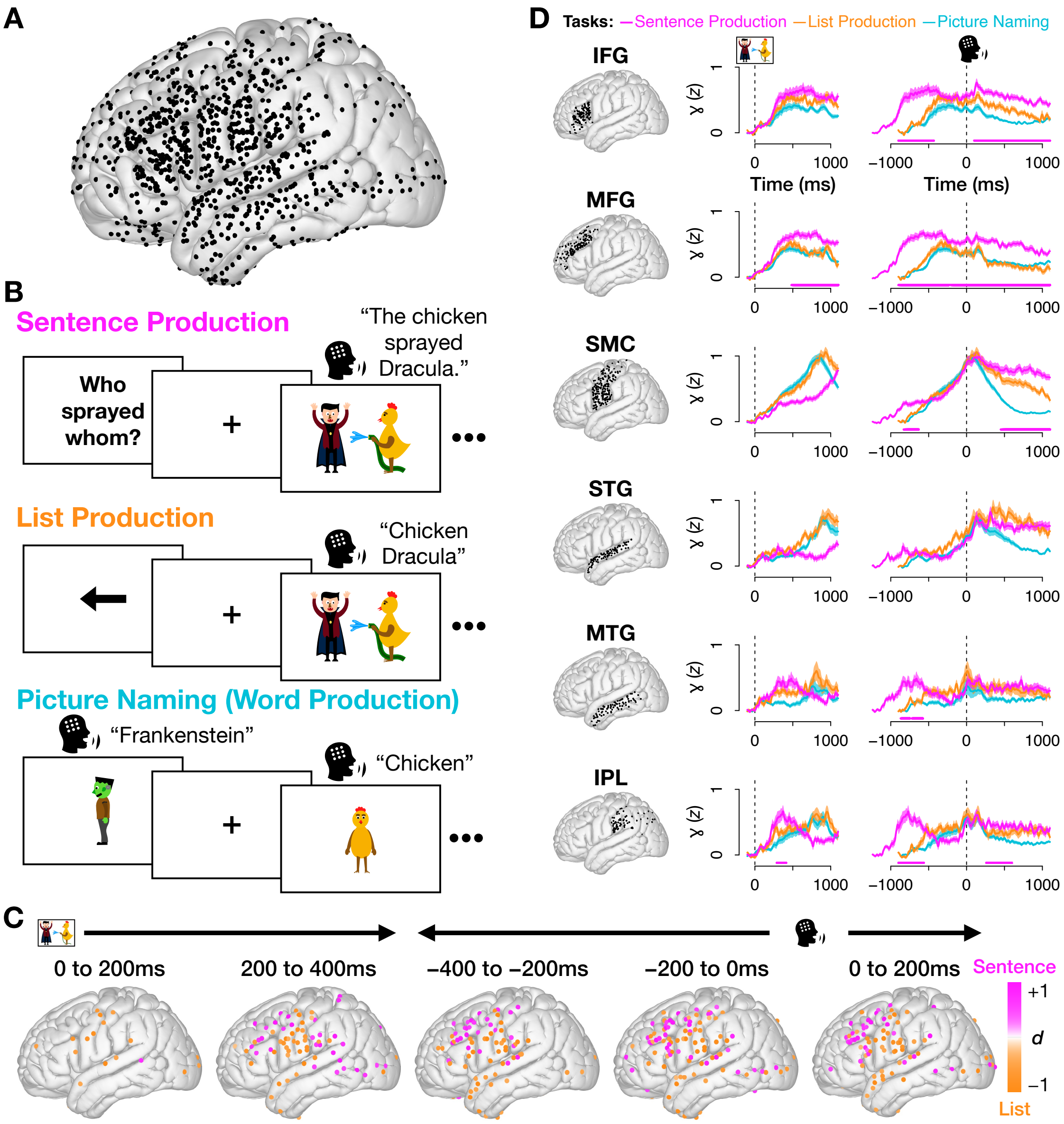


**Figure S2**: **Sentence–list high gamma effects are robust to temporal warping of the planning period.** Here we replicate Figure 1 using warped data. (A) Our coverage (unchanged). (B) Experimental paradigm (unchanged). (N.B.: The cartoon figures here and throughout are not the exact images used in the experimental stimuli, but are visually similar. They were created by the authors (*149*) for publication.) Results are qualitatively identical: (C) the spatial distribution of electrodes with significant differences between sentences and lists over time was nearly identical and (D) both identify IFG, MFG, MTG, and IPL as having significantly higher sentence than list activity during the planning period.

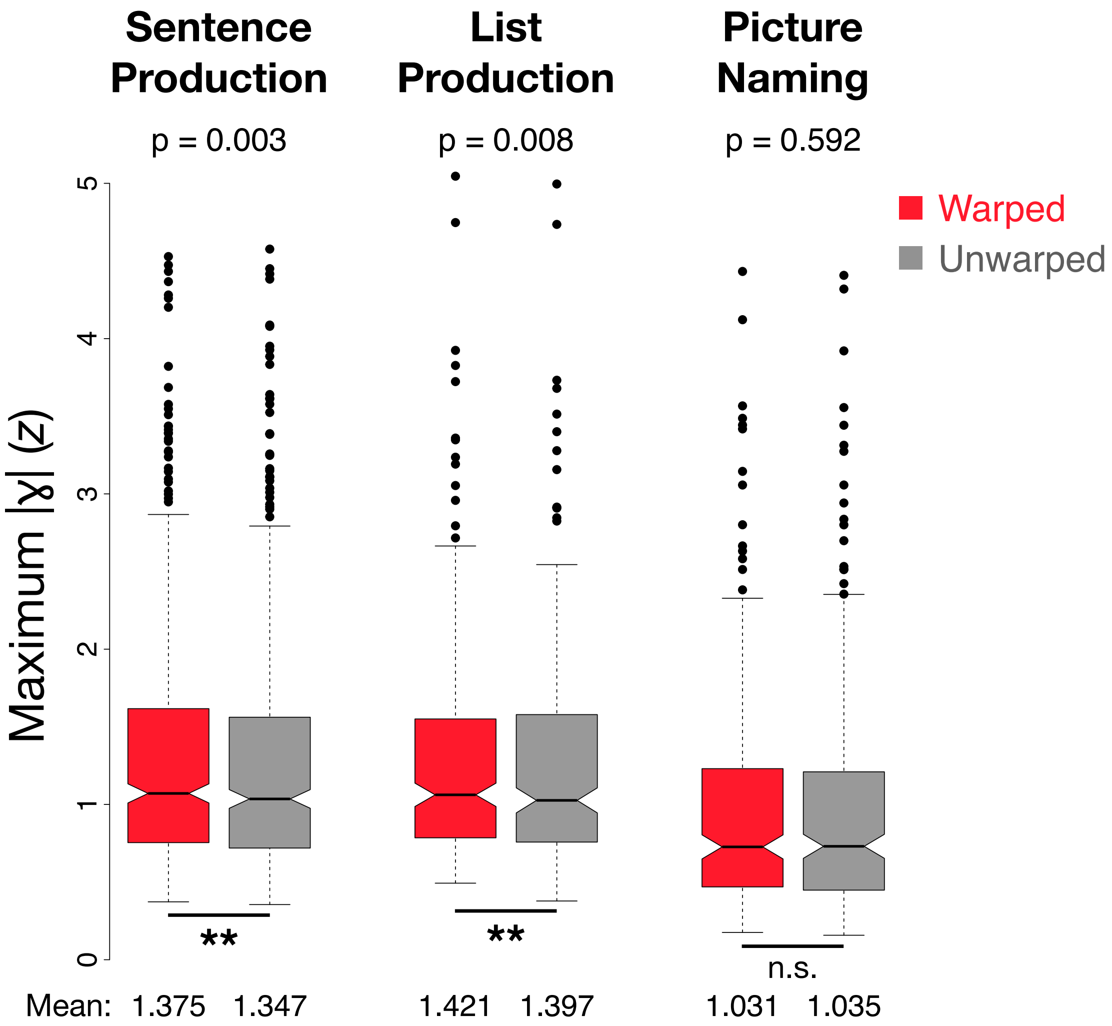


**Figure S3**: **Temporal warping improves alignment of high gamma during sentence and list planning.** Peak high gamma activity for each electrode (dot) in the warped (red) and unwarped (grey) data, by task. Warping resulted in significantly higher peaks for sentence production ($p$ = 0.003, Wilcoxon signed rank test) and list production ($p$ = 0.008, Wilcoxon signed rank test), evidence that it successfully improved the temporal alignment of trials (*100*). No effect was found for picture naming ($p$ = 0.592).

#### Analyses of Beta Activity

Supplementary Fig. S4 replicates the Figure 1 analyses but using beta activity (12-30 Hz). Beta is another important frequency band in cognition (*91, 92*). In Supplementary Fig. S5, we test the hypothesis that beta activity tracks structural differences more clearly than high gamma. We present three replications of the analysis in Fig. 3A (copied in Supplementary Fig. S5A). Analyses appear in a $2 \times 2$ factorial design: regressing either beta or high gamma activity over RSIs, where RSIs were derived from RSA performed on either high gamma (left) or beta (right) activity. Across all four sets of results, the only significant finding is that reported in the main text: a positive relationship between mean sentence high gamma and (sub)lexical signal in the high gamma trial activity ($p < 0.001$, linear regression).

To determine whether representational encoding in beta activity might be anatomically localized more than in high gamma, we replicated the analyses in Fig. 2 using beta. We analyzed the resulting "beta RSIs" by region of interest (Supplementary Fig. S6A). There was significant evidence for sensitivity to structure in SMC and, as in the high gamma data, IFG ($p < 0.05$ for 100 ms, permutation test), but unlike high gamma there was no evidence that beta activity was sensitive to the active/passive contrast in MFG. We also plotted electrodes with significant beta RSIs on cortex (Supplementary Fig. S6B), revealing distributed networks similar to those observed in high gamma.

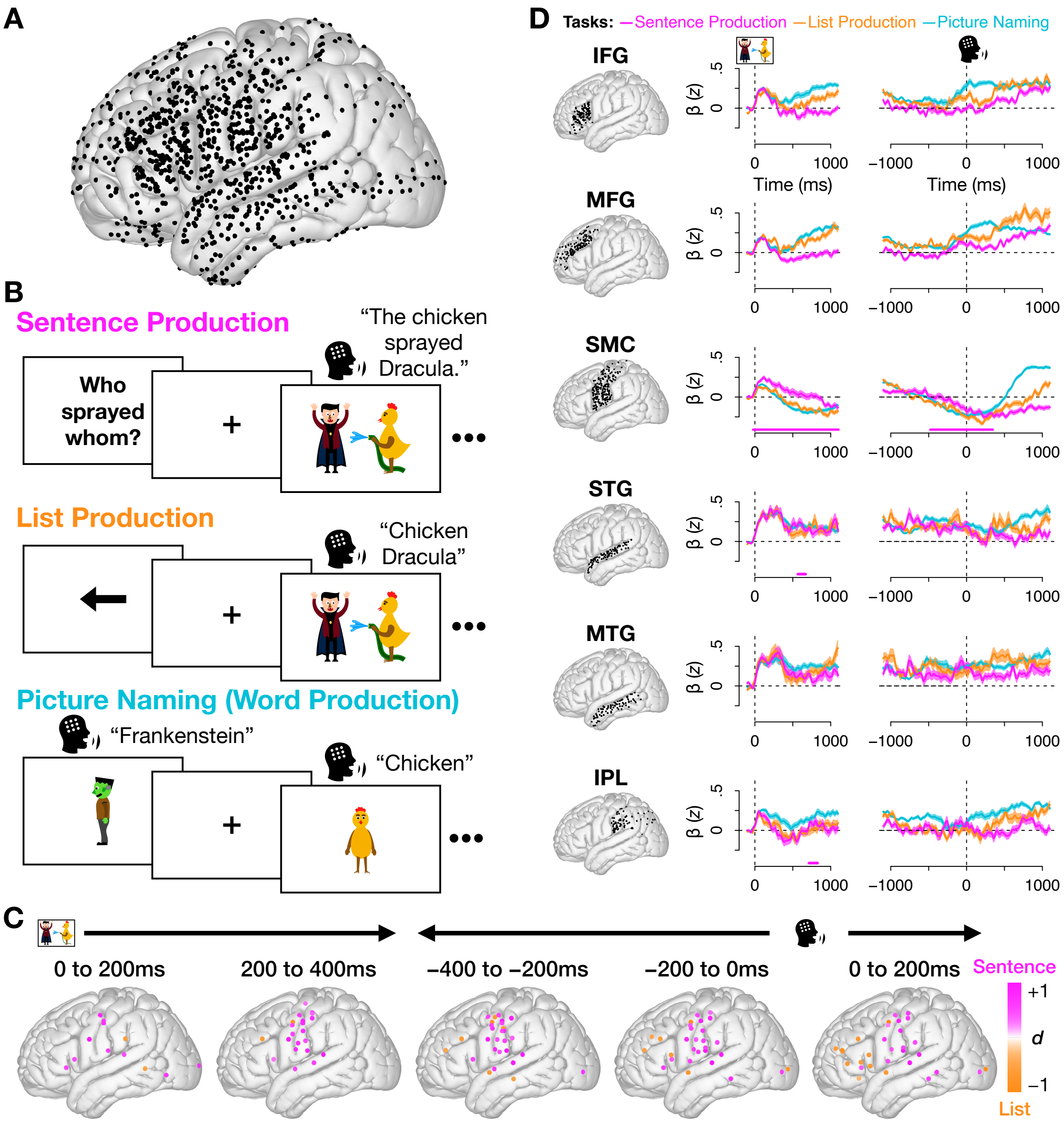


**Figure S4**: **Beta-band activity shows suppression corresponding to high gamma increases.** Replication of Figure 1 using beta activity (12-30 Hz; unwarped data). (C) The distribution of electrodes that are significantly greater for sentences than lists was largely reversed from the high gamma activity. This likely reflects a well-documented phenomenon where activity in high gamma is often coupled with beta suppression, particularly in sensorimotor regions, leading to effects in the reverse direction in beta (*150, 151*). (D) SMC, STG, and IPL show significantly higher beta activity for sentences than lists, possibly also reflecting beta suppression corresponding to increases in high gamma during speech and auditory feedback.

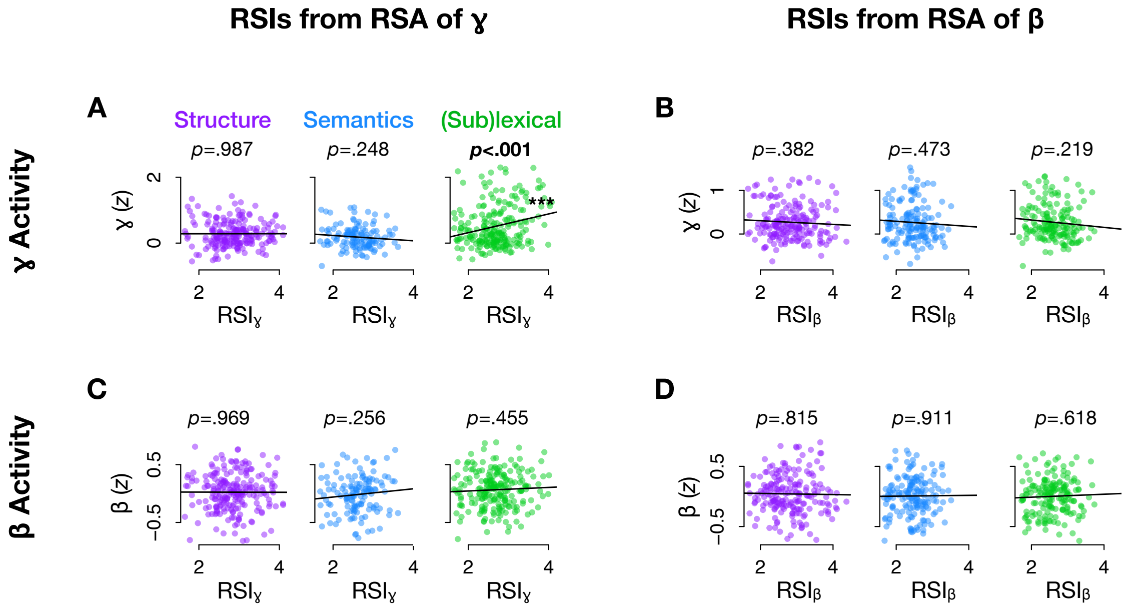


**Figure S5**: **Across beta and high gamma, mean activity tracks (sub)lexical but not higher-order linguistic processing.** Four attempts at uncovering a relationship between mean neural activity during sentence production and linguistic processes. Only one significant relationship was found across all analyses: mean high gamma activity was positively related to high gamma (sub)lexical processing ($p < 0.001$). (A) Mean high gamma activity vs. linguistic RSIs encoded in high gamma trial activity (panel is identical to Fig. 3A). (B) Mean high gamma activity vs. linguistic RSIs encoded in beta trial activity. (C) Mean beta activity vs. linguistic RSIs encoded in high gamma trial activity. (D) Mean beta activity vs. linguistic RSIs encoded in beta trial activity.

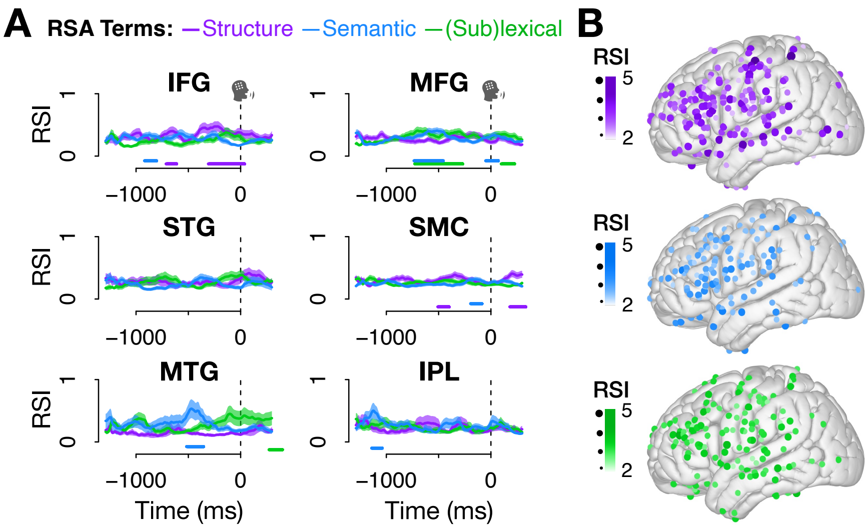


**Figure S6**: **Distributed beta-band sensitivity to linguistic information parallels high gamma RSIs.** Distribution of RSIs calculated from RSA on beta activity (see parallels for high gamma in Fig. 2). (A) Mean and standard error by region shows significant structure effects in IFG and SMC ($p < 0.05$ for 100 ms, permutation test), but there was no evidence for sensitivity to structure in MFG as there was in the high gamma activity. (B) Significant electrodes ($p < 0.05$ for 100 ms; one-tailed permutation test) per RSI again show a broadly distributed pattern for all three RSIs, as in high gamma.

#### Anatomy of RSIs and their overlap

Figure S7 shows the distribution of electrodes which were significant for each RSI (and each combination of RSIs) during the planning period.

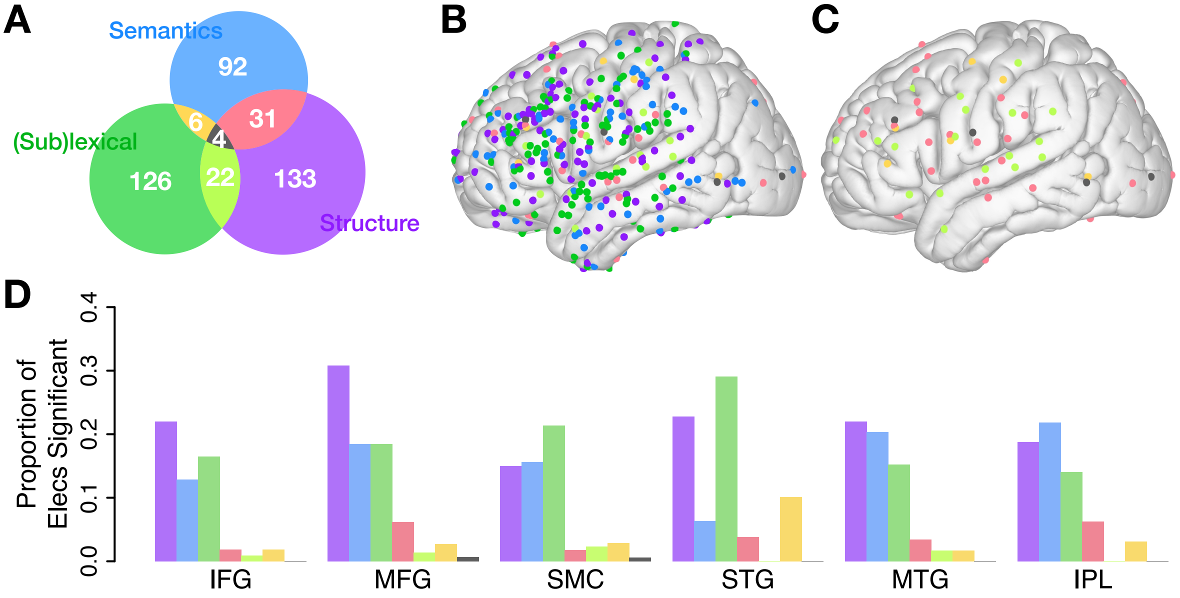


**Figure S7**: **Anatomical distributions of RSIs.** (A) The number of electrodes significant for each RSI during the planning period (a replication of Fig. 2E to serve as a color key for the remaining panels in this figure). (B) All electrodes significant for one or more RSI. Colors correspond to the Venn diagram in Panel A, indicating which RSI(s) each electrode was significant for. (C) The distribution of overlapping structural, event semantic, and/or (sub)lexical information (i.e., just electrodes significant for two or three RSIs). (See Fig. 4A-B for the distribution of individual RSIs.) (D) The proportion of analyzed electrodes in each ROI which were significant for each RSI (and combinations therein).
